# TREM2 expression level is critical for microglial state, metabolic capacity and efficacy of TREM2 agonism

**DOI:** 10.1101/2024.07.18.604115

**Authors:** Astrid F Feiten, Kilian Dahm, Bettina van Lengerich, Jung H Suh, Anika Reifschneider, Benedikt Wefers, Laura M Bartos, Karin Wind-Mark, Kai Schlepckow, Thomas Ulas, Elena De-Domenico, Matthias Becker, Igor Khalin, Sonnet S. Davis, Wolfgang Wurst, Nikolaus Plesnila, Jonas J Neher, Matthias Brendel, Joseph W Lewcock, Gilbert Di Paolo, Anja Capell, Kathryn M Monroe, Joachim L Schultze, Christian Haass

## Abstract

Triggering receptor expressed on myeloid cells 2 (TREM2) is a central regulator of microglial activity and sequence variants are major risk factors for late onset Alzheimer’s disease (LOAD). To better understand the molecular and functional changes associated with TREM2 signalling, we generated a TREM2 reporter mouse model and observed a gradual upregulation of reporter expression with increasing plaque proximity. Isolated microglia were sorted based on reporter expression and their transcriptomic profiles acquired in both wildtype and APP transgenic animals, allowing us to disentangle TREM2 versus pathology-specific effects. Bulk RNA-sequencing highlighted TREM2 level-dependent changes in major immunometabolic pathways, with enrichment of genes in oxidative phosphorylation and cholesterol metabolism in microglia with increased TREM2 expression. To confirm these findings, we next analysed uptake of fluorodeoxyglucose (FDG) and examined metabolomic and lipidomic profiles. Again, independent of Aβ pathology, TREM2 expression correlated with uptake of FDG as well as increased cellular redox, energetics, and cholesterol homeostasis. Finally, we performed chronic treatment with a brain penetrant TREM2 agonist and identified a window of TREM2 expression where microglia are most responsive. Thus, our data provide novel insights into TREM2-mediated regulation of microglial metabolic function and informs current efforts to bring TREM2 agonists into clinical application.

## Introduction

Although the original descriptions of Alzheimer’s disease (AD) pathology already suggested glial pathology, its contribution to disease progression remained largely controversial until genome wide association studies (GWAS) identified risk loci for late onset AD (LOAD) expressed predominantly in microglia^1^. Now microglia are known to form a barrier around amyloid plaques^2–5^. These plaque-associated microglia are phagocytic and mediate removal of aggregated amyloid β-peptide (Aβ)^6^. Furthermore, this barrier restricts the amount of toxic Aβ oligomers that can escape from the plaque, and thereby may limit neuritic dystrophy and neurotoxicity^3,4^. In order to perform these protective functions, microglia must change their activation state^7^. The major regulator of the transition from homeostatic microglia to disease-associated microglia (DAM) is the triggering receptor expressed on myeloid cells 2 (TREM2)^7–9^. Heterozygous sequence variants of this transmembrane receptor have been associated with LOAD^10,11^ and frontotemporal dementia (FTD)^12,13^, while homozygous mutations in TREM2 or its signalling adaptor DAP12 (encoded by the gene *TYROBP*) are causative for Nasu-Hakola disease (NHD), an early onset dementia with bone cysts^14^. Within the brain, TREM2 is predominantly expressed in microglia^15^, where it forms a complex with DAP12 to initiate downstream signalling^16,17^. Activation of TREM2 has been associated with functions such as phagocytosis^8,18,19^, synaptic pruning^20^, and controlling the inflammatory state through cytokine and chemokine release^18,21^. It has also been suggested that TREM2 can regulate tau kinases^22^, glucose uptake^23–25^, and microglial metabolism^23,25–27^, linking it to core AD pathologies. Furthermore, AD patients with underlying *TREM2* loss-of-function variants and *Trem2* knockout (KO) mice show a reduction of microglial clustering around amyloid plaques^5^ and a downregulation of mammalian target of rapamycin (mTOR), which plays an important role in cellular metabolism^26^. Microglial TREM2 has been shown to be vital in clearing myelin debris as the induction of a DAM state seems to be required to efficiently clear cholesteryl esters derived from myelin cholesterol^28,29^. In the periphery, TREM2 is a master regulator of lipid-associated macrophages (LAM)^30^. In adipose tissue, loss of TREM2 leads to systemic hypercholesterolemia, inflammation and glucose intolerance, suggesting a critical function of TREM2 in energy metabolism and lipid catabolism^30^.

Recently, different agonistic antibodies targeting TREM2 have been explored as treatment options for LOAD^31,32^. Agonistic antibodies stimulate microglial proliferation, mitochondrial metabolism and glucose uptake^25,32,33^ and some of them reduce amyloid plaque load^34^. First clinical trials with TREM2 agonists are currently in early phases^32^. However, for the design of disease modifying trials and to achieve the best treatment efficacy, clinicians need to be informed about the state of microglia activation, which allows optimal stimulation of TREM2-dependent protective functions.

Here, we generated a TREM2 reporter mouse model to investigate how TREM2 expression levels in amyloid-burdened and healthy mice affect metabolic functions of microglia. We isolated microglia and performed cell sorting based on the expression of the fluorescent TREM2 reporter, which allowed for analyses of subpopulations of microglia expressing low, mid, and high TREM2 from the same mouse. Notably, we found that the amount of TREM2 directly relates to glucose uptake levels and is an important regulator of microglia metabolism, independent of disease context. The metabolic and lipidomic profile of high TREM2 expressing microglia was shifted towards increased phagocytosis, increased energetic and metabolic capacity, and decreased cholesterol, suggesting a protective microglial change induced by robust TREM2 activation. Moreover, treatment with an agonistic TREM2 antibody revealed mid TREM2 expressing cells to be the most responsive. These findings confirm TREM2 as a crucial and dose-dependent regulator of microglial metabolism and have important implications for the design of upcoming clinical trials.

## Results

### Generation of a TREM2 reporter mouse reveals gradual upregulation of TREM2

We generated a TREM2 reporter mouse to investigate how differences in TREM2 expression and its functional consequences correlate with amyloid plaque pathology. We used the fluorophore mKate2^35^ as a reporter, separated from *Trem2* by a P2A spacer to express both proteins individually under the control of the same promotor^36^. To ensure that the mKate2 reporter protein is retained within the cell, we attached the endoplasmic reticulum (ER)-retention sequence ‘KDEL’^37^ to its C-terminus (**Fig. 1A**). We first confirmed the expression of both proteins TREM2 and mKate2 from the polycistronic cDNA construct in HeLa cells (**Fig. 1B**). mKate2 was expressed evenly throughout the endoplasmic reticulum and clearly separated from cell surface TREM2 staining (**Fig. 1B**). Using CRISPR/Cas9, we next introduced the P2A-mKate2-KDEL cassette 3’ of exon 5 into the endogenous mouse *Trem2* locus (**Fig. 1C**) and further analysed homozygous (*Trem2-mKate2^KI/KI^*, hereafter *Trem2^KI/KI^*), heterozygous (*Trem2-mKate2^KI/wt^*, hereafter *Trem2^KI/w^*^t^), and wild type (*Trem2-mKate2^wt/wt^*, hereafter *Trem2^wt/wt^*) mice. Co-staining of mKate2 and the microglial marker Iba1 showed that mKate2 expression localised to microglia (**Fig. 1D**). Insertion of the mKate2 cassette did not significantly alter total amounts of *Trem2* mRNA (**Extended Data Fig. 1A**). As expected, a gene dose dependent increase of *mKate2* mRNA was observed (**Extended Data Fig. 1B**), with a corresponding decrease of the unmodified (wt) *Trem2* mRNA, i.e. mRNA lacking the silent mutations introduced during generation of the reporter line (**Extended Data Fig. 1C**). Next, we confirmed microglial activation as a proxy for TREM2 receptor function in the reporter mice using a well-established model for controlled cortical impact (CCI; **Extended Data Fig. 1D**)^38^. Traumatic brain injury was shown to trigger microgliosis associated with an upregulation of DAM markers^39,40^. 72h post injury, *Trem2^wt/wt^* as well as *Trem2^KI/wt^* mice showed an upregulation of total *Trem2* mRNA on the ipsilateral site compared to naïve mice (**Extended Data Fig. 1E**). As expected, in *Trem2^KI/wt^*mice we found an upregulation of both the *mKate2* reporter RNA (**Fig. S1F**) and the unmodified wt *Trem2* RNA (**Extended Data Fig. 1G**). The DAM markers *Clec7a*, and *Cd68*^7^ as well as *Grn* were increased to a similar extent in *Trem2^wt/wt^*and *Trem2^KI/wt^* mice (**Extended Data Fig. 1H**), suggesting that microglial transition to a DAM state is not altered by the genomic insertion of the reporter.

**Figure 1.**
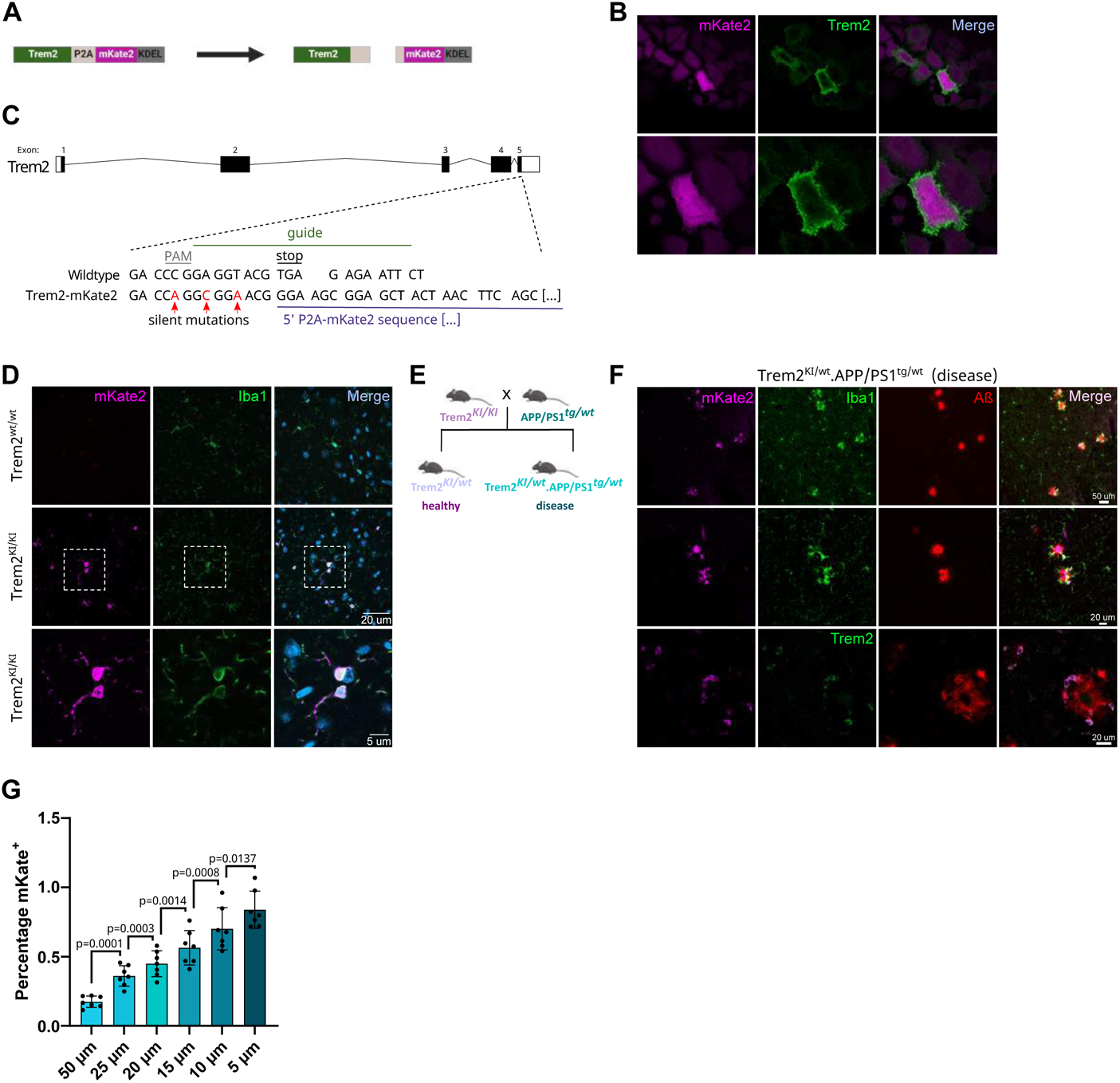
A novel TREM2 reporter mouse reveals gradual upregulation of TREM2 in microglia. (A) Schematic of the reporter construct and the proteins expressed in-vitro. (B) Representative immunofluorescent images of transfected HeLa cells. TREM2 staining is displayed in green and mKate2 in magenta. (C) CRISPR strategy to create the *Trem2-mKate2* reporter mice. (D) Representative immunofluorescent images of *Trem2-mKate2* mice (wt and homozygous KI) showing positive labelling of mKate2 only in knock in animals (magenta). Dashed squares represent the area of the zoom in in the bottom row. Iba1 staining in green and DAPI in blue. Scale bar 20 μm and 5 μm respectively. (E) Breeding scheme used to create the experimental mice for the *Trem2-mKate2.APP/PS1* strain. Only heterozygous reporter mice were used throughout the study. Non-transgenic littermates are referred to as ‘healthy’ and APP/PS1 transgenic mice are hereafter called ‘disease’. (F) Representative immunofluorescent image of disease mice (4 months). mKate2 is displayed in magenta, Iba1 positive microglia or Trem2 in green and amyloid beta (Aβ) plaques in red. Scale bar 50 μm and 20 μm respectively. (G) Percentage of mKate2 positive signal with decreasing distance to a plaque. One data point represents averages of multiple images per mouse, n = 7, age = 4 months, error bars represent standard deviation.

To examine the dynamics of TREM2 expression in the presence or absence of amyloid plaques, we crossed the reporter line with the APP/PS1 mouse^41^, a model of amyloidosis, to generate *Trem2-mKate2^KI/wt^.APP/PS1^tg/wt^*(hereafter called “disease”) mice and compared them to *Trem2-mKate2^KI/wt^* (hereafter called “healthy”) mice (**Fig. 1E**). Histological analyses of 4-month-old mice revealed detectable mKate2 expression predominantly but not exclusively in microglia in close proximity to plaques (**Fig. 1F**). Calculating the percentage of mKate2 positive staining around plaques showed a gradual upregulation of the signal with increasing plaque proximity (**Fig. 1G**), suggesting the existence of microglia subpopulations with varying expression levels of TREM2.

### TREM2 levels determine expression of functionally distinct microglial gene modules

To further characterize potential microglial subpopulations, we isolated total brain microglia from 9-month-old animals and performed fluorescence activated cell sorting (FACS) to separate microglia based on the mean fluorescence intensity of the mKate2 reporter as a proxy for TREM2 expression. We defined mKate2 low-, mid- and high expressing subpopulations in disease animals, and low- and mid mKate2 expressing microglial subpopulations in healthy animals (**Fig. 2A**). As expected, high mKate2 expressing subpopulations were almost absent in healthy animals (**Fig. 2A**), although the relatively wide range of TREM2 expression observed in these animals suggests microglial subpopulations expressing varying amounts of TREM2 exist even under physiological conditions (**Fig. 2A**). This is consistent with an age dependent increase of the TSPO-PET signal and the formation of microglia nodules at sites of small myelin damage previously observed in aged wildtype mice^23^.

**Figure 2.**
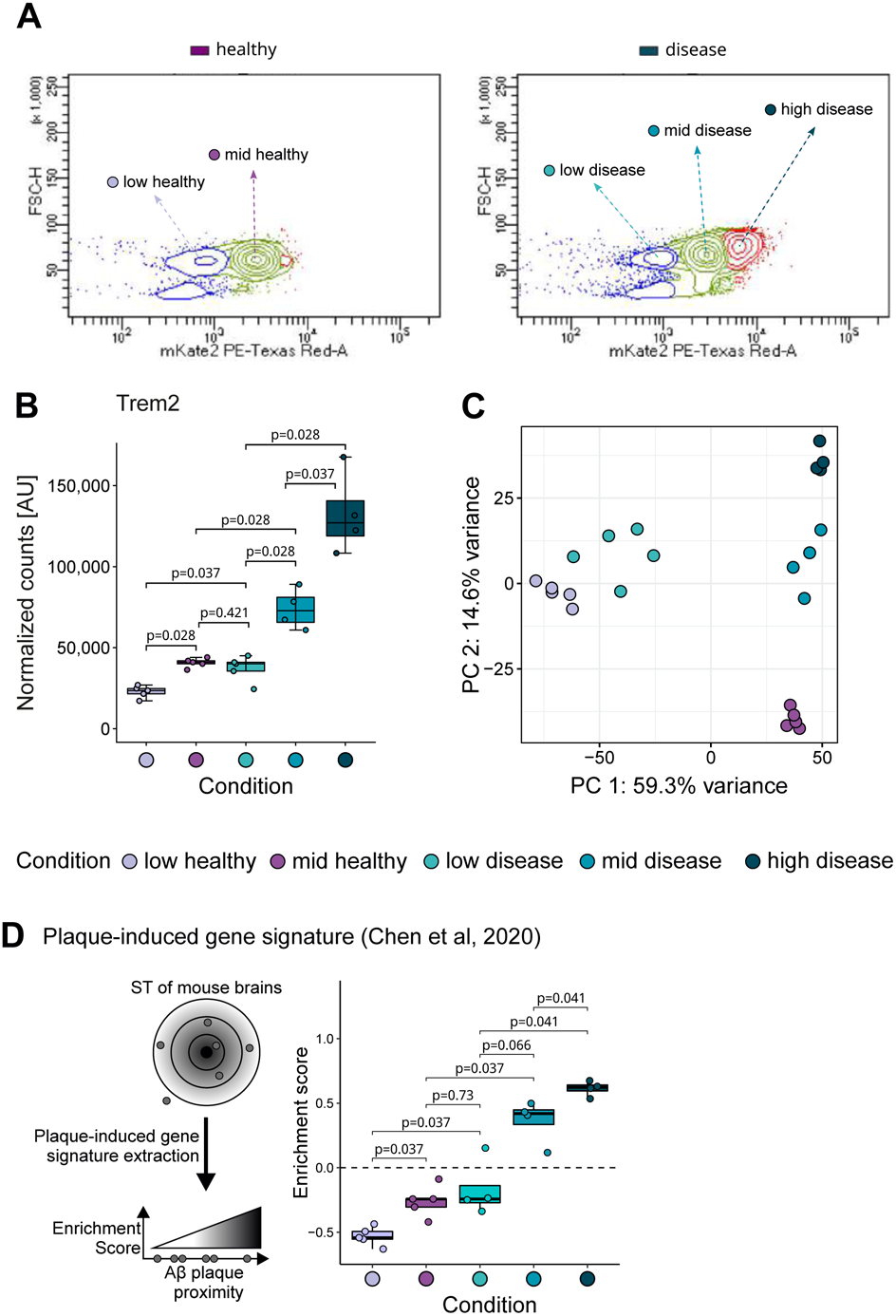
Microglia upregulate plaque-induced and disease-associated signature in a TREM2-dependent manner. (A) Example contour plots showing the rational for the FACS gating strategy in healthy and disease mice based on the amount of mKate2 positivity, as a proxy for TREM2 expression. Subpopulations were sorted and named based on the amount of mKate2 present. (B) Boxplot of normalized TREM2 expression coloured by condition. Boxplots show the 25%, 50% (median) and 75% percentile, whiskers denote 1.5 times the interquartile range (applies to all following boxplots). For all RNAseq data n = 4-5 mice per condition, age = 9 months (C) PCA of top 5,000 most variable genes coloured by condition. (D) Boxplot of gene set variation analysis (GSVA) enriching for plaque induced gene signature from Chen et al. coloured by condition. Error bars represent the standard deviation, p-values calculated with Wilcoxon rank sum test with Benjamini-Hochberg adjustment (B,D)

To characterize the biological relevance of microglial subpopulations stratified by TREM2 expression and disease pathology, we subjected sorted mKate2 low, mid, and high microglia to bulk RNA-seq analysis (workflow **Extended Data Fig. 2A**). Transcriptomic analysis confirmed that the microglial subpopulations identified via flow cytometry expressed different levels of *Trem2* transcripts as *Trem2* mRNA levels across microglial subpopulations showed a gradual increase (**Fig. 2B)**. To explore whether microglia with different TREM2 levels were characterized by distinct transcriptional programs, we performed PCA on the 5,000 most variable genes (**Fig. 2C**). Assessment of low-dimensional embedding revealed distinct clusters for each group. PC1 explained almost 60% of the variance in the data and separated the clusters based on TREM2 levels, demonstrating a major impact of TREM2 expression on global transcriptomic responses. PC2 further split the clusters according to disease state (**Fig. 2C)**.

Subsequently, we determined to what degree these TREM2 driven transcriptomic differences related to amyloid plaque proximity by performing a gene set variation analysis (GSVA) of a plaque-induced gene (PIG) signature previously defined based on spatial transcriptomics and single-cell data (**Fig. 2D**)^42^. We observed enrichment of the PIG signature only in microglia from mid and high mKate2 disease conditions with a gradually increasing enrichment score, while the PIG signature was not increased in the mid mKate2 healthy and low mKate2 disease population, whereas the low mKate2 healthy subpopulation was even negatively enriched (**Fig. 2D**). To assess the ranking of the PIGs on a population level, a gene set enrichment analysis (GSEA) was performed, corroborating positive enrichment of signature genes in mid and high mKate2 disease populations and negative enrichment in the low mKate2 healthy population. (**Extended Data Fig. 2B**). On the other hand, within the mid mKate2 healthy and low mKate2 disease microglia the PIGs ranking resembled the enrichment pattern of random signatures on population and sample level (**Extended Data Fig. 2C – 2E**). To further validate the microglial subpopulations we performed cell state deconvolution based on linear support vector regression (LSVR) utilizing reference single-cell RNA-seq data sets previously established in two independent AD models (**Extended Data Fig. 2F** homeostatic vs DAM^7^, **Extended Data Fig. 2G** Methoxy XO4 positive vs negative)^7,43^. The comparisons with previously published microglial phenotypes (PIG, DAM, XO4+) confirm that these signatures develop in response to Aβ pathology and demonstrate that these expression programs increase with TREM2 expression levels.

Our data indicated the presence of disease-independent TREM2-driven as well as TREM2-independent disease-driven transcriptional alterations. To delineate these different gene expression programs, we compared low mKate2 disease microglia with low mKate2 healthy microglia, and mid mKate2 disease microglia with mid mKate2 healthy microglia (**Fig. 3A**), and intersected the differentially expressed genes (DEGs) of both comparisons, which resulted in 106 DEGs (from here on defined as AD-related DEGs) (**Extended Data Table 1**). Heatmap visualization (including high disease) clearly shows the differences between healthy and disease for both groups even though the expression levels of the majority of these genes gradually increased with increasing TREM2 levels (**Fig. 3B**). Next, we defined genes directly related to TREM2 levels, by performing comparisons between groups with different TREM2 levels but matching health status (mid mKate2 healthy vs low mKate2 healthy, mid mKate2 disease vs low mKate2 disease, high mKate2 disease vs low mKate2 disease and high mKate2 disease vs mid mKate2 disease) (**Fig. 3C**). Intersection of those comparisons resulted in 65 DEGs (from here on defined as TREM2-related DEGs) (**Fig. 3D, Extended Data Table 1**). As expected, GSVAs of both the AD-related and TREM2-related DEGs significantly split the microglia according to their health status and TREM2 level, respectively (**Extended Data Fig. 3A, 3B**)^43^. Moreover, among the DEGs, the DAM associated genes *Clec7a* and *Apoe*^7^ were also identified (**Fig. 3E**). Like *Trem2* **(Fig. 2B**), both genes showed a gradual increase in expression in the disease condition from low to high mKate2; however, in the healthy condition a significant decrease in expression from low to mid mKate2 was observed, indicative of a differential transcriptional program associated with TREM2 levels in healthy animals. The TREM2 downstream signalling genes *Hcst* (transcript of Dap10) and *Tyrobp* (transcript of Dap12), known to be involved in *Syk*-independent^44^ and *Syk*-dependent *Trem2*-signaling^44,45^, respectively, had a similar expression distribution compared to *Trem2* across conditions (**Fig. 3F**), while *Syk* itself, showed a decrease in expression from low mKate2 healthy to the high mKate2 disease condition, possibly indicating a negative feedback loop. Interestingly, *Mtor*, a major metabolism and energy sensor and regulator of pathways downstream of TREM2, was equally expressed across all conditions (**Fig. 3F**)^26,44^.

**Figure 3.**
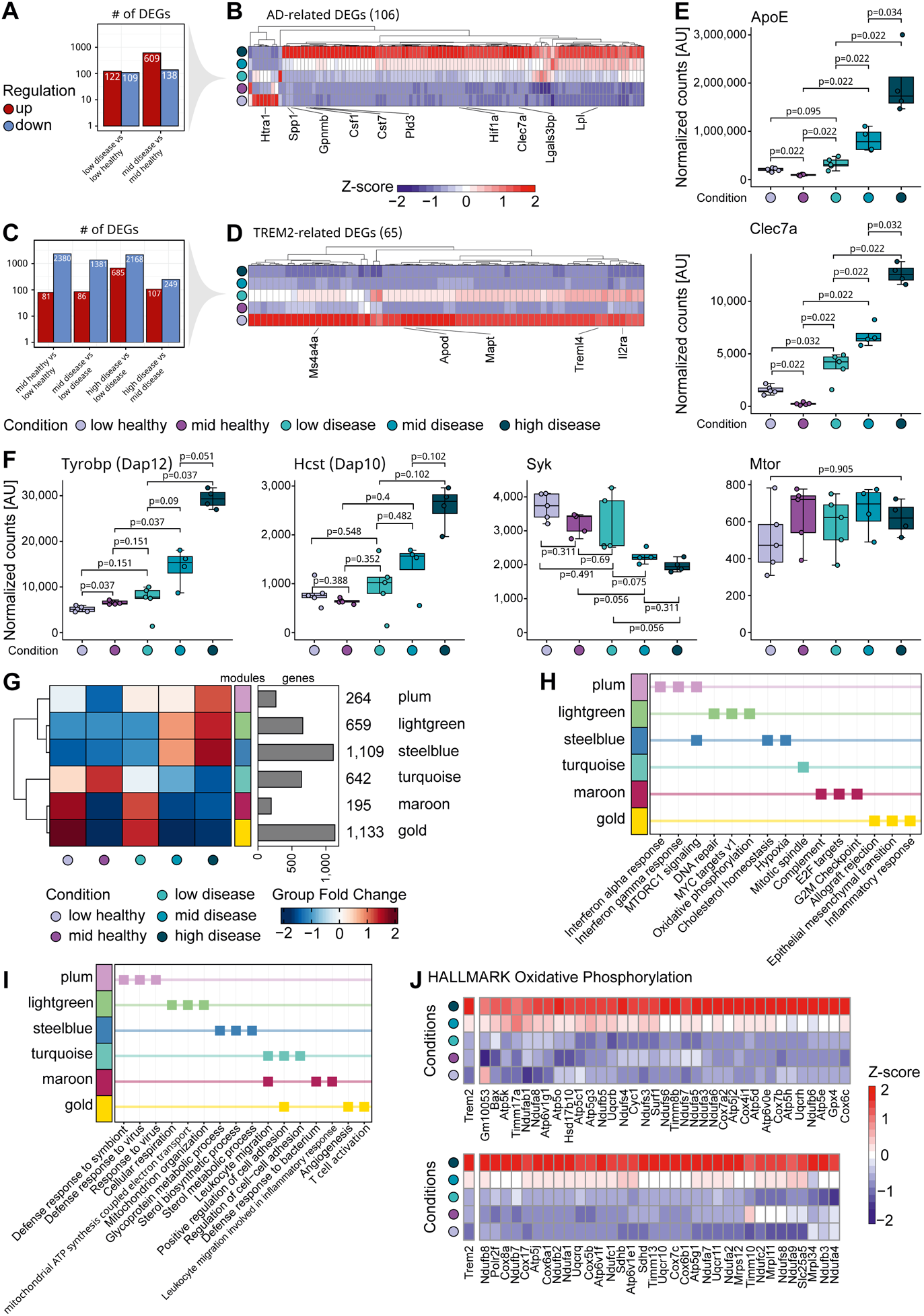
Functionally distinct microglia gene modules are differentially driven by TREM2 expression level. (A) Bar plot of DEGs between healthy and disease mice with matching TREM2 level coloured according to the mode of regulation (red: up; blue: down). (B) Clustered heatmap of AD-related DEGs from comparisons in A, additionally high disease was plotted for comparison. Each DEG is coloured by its normalized, z-scored gene expression. Selected DEGs are highlighted. (C) Bar plot of DEGs between different TREM2 levels with matching health status coloured according to the mode of regulation. (D) Clustered heatmap of TREM2-related DEGs from comparisons in C. Each DEG is coloured by its normalized, z-scored gene expression. Selected DEGs were highlighted. (E) Boxplot of normalized gene expression of *ApoE* and *Clec7a* coloured by condition. (F) Boxplot of normalized gene expression of *Tyrobp* (transcript of Dap12), *Hcst* (transcript of Dap10), *Syk* and *Mtor* coloured by condition. (G) Clustered heatmap of gene co-expression network modules split by condition and coloured by group fold change (left), and bar plot of number of genes included in the respective module (right). (H) Overview of Hallmark enrichment of gene modules displaying the top 3 associated terms per gene module based on the adjusted p-value. Squares are coloured by module. (I) Overview of GO enrichment of gene modules displaying the top 3 associated terms per gene module based on the adjusted p-value. Squares are coloured by module. (J) Heatmap of genes associated with the Hallmark term ‘Oxidative phosphorylation’ in the light green module coloured by normalized and z-scored expression and ordered by condition. p-values calculated with Wilcoxon rank sum test with Benjamini-Hochberg adjustment (E,F)

To further dissect transcriptional programs related to TREM2 function versus disease pathology, we performed gene co-expression network analysis. The resulting network was defined by six major gene modules, containing 4,002 genes. When plotting condition-specific expression patterns in a heatmap, significant differences were revealed (**Fig. 3G**). The modules plum, lightgreen and steelblue showed enrichment in the high mKate2 disease followed by the mid mKate2 disease conditions. Interestingly, the plum module was enriched for type I and II interferon response genes (*Ifitm3*, *Stat1*, *Irf7*, *Irf9*; **Fig. 3H**), which were recently identified to be uniquely upregulated after treatment with a TREM2 agonist^25^. The lightgreen and steelblue modules were associated with an increased expression of genes related to metabolic activity, namely ‘oxidative phosphorylation’ (*Atp5j, Cox5b, Uqcrq*) in the lightgreen module and ‘sterol metabolic processes’ (*Apoe*, *Cebpa*) and ‘cholesterol homeostasis’ (*Cd9*, *Lpl*, *Tmem97*) in the steelblue module (**Fig. 3H, 3I**). Further, a closer inspection of the genes involved in the hallmark ‘oxidative phosphorylation’ revealed a gene expression pattern matching the *Trem2* expression level of the respective microglial condition (**Fig. 3J**). The gold and maroon modules consisted of genes with high expression levels in microglia with low TREM2 levels, independent of disease state (**Fig. 3G**) and were functionally related to genes involved in ‘G2M Checkpoint’ (*Mki67*), suggesting they have proliferation potential together with an enrichment of genes required for ‘Inflammatory response’ (*Cd40*, *Tnfaip6*) (**Fig. 3H**) and ‘T-cell activation’ (*Cd74*, *Clec4a1*) (**Fig. 3I**). Taken together, the transcriptome analysis of microglia subpopulations based on TREM2 expression levels in healthy and diseased mice revealed both TREM2-as well as disease-related gene modules, indicating differential transcriptional responses as a result of TREM2 expression that are further shaped by exposure to amyloid plaques.

### Gradual upregulation of TREM2 in plaque proximity is associated with increased glucose uptake

We previously observed that microglia alter their energy metabolism in the presence of a pathological challenge^46^. In line with this finding, we demonstrated that glucose uptake in microglia is TREM2-dependent and modulated by activation state^24^. Here, our bulk RNAseq data revealed that genes involved in oxidative phosphorylation were altered in the subpopulations with different amounts of TREM2. To directly examine if and how TREM2 levels in microglial subpopulations could influence energy metabolism, we injected 13-14-month-old mice with 45 ± 3 MBq [^18^F]FDG. CD11b-enriched microglia were sorted into low, mid and high expressing subpopulations as defined by mKate2 levels and radioactivity was measured (**Fig. 4A**)^47^. The proportion of mKate2 low microglia was roughly 25% in both healthy and disease mice (**Fig. 4B**) while the mKate2 high subpopulation was virtually absent in healthy mice. Glucose uptake increased with mKate2 expression although it reached significance only in disease mice (**Fig. 4B**). The unexpected similar levels of glucose uptake by mid mKate2 healthy and mid/high mKate2 disease microglia may be due to the downregulation of S*yk* signalling in the mid/high mKate2 disease microglia (see Fig. 3F) or indicate that the amount of TREM2 expressed by mid mKate2 healthy microglia might be sufficient to trigger maximal glucose uptake.

**Figure 4.**
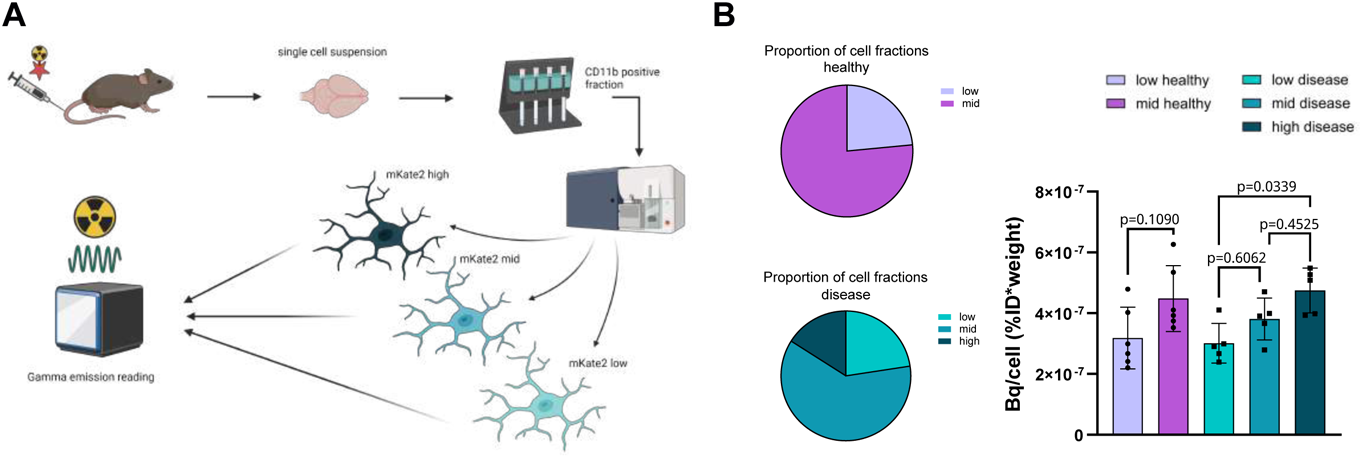
Glucose uptake is associated with TREM2 upregulation. (A) Schematic showing the workflow of scRadiotracing following radiolabelled FDG injection, here in 14 months old *Trem2-mKate2.APP/PS1* mice. (B) Quantifications of cells sorted after radiolabelled FDG injection. Left: proportion of subpopulations; right: quantification of FDG taken up by each cell. N = 5-6, age = 14 months.

### Distinct metabolic profiles are associated with TREM2 expression level

Since the above findings suggest that increased glucose uptake and gene expression related to metabolic changes correlate with increased TREM2 expression in healthy and diseased mice, we sought to investigating and validate whether TREM2 regulates the microglial metabolic state in the presence and absence of amyloid pathology. To test this, metabolomic and lipidomic analyses were conducted in sorted healthy and disease microglial subpopulations. To identify major trends in metabolic processes according to diseases status or TREM2 expression levels, principal component analysis (PCA) was performed to reduce the dimensionality of the data. Consistent with glucose uptake, liquid chromatography-mass spectrometry (LC-MS) findings suggest that TREM2 levels, rather than amyloid pathology, drive component stratification (**Fig. 5A**). As such, TREM2 expression tracked more closely with patterns of metabolic variations along the PC1 axis than disease status (**Fig 5B**). PC1 scores were positively contributed by cellular energetic substrates, such as adenosine, S-adenosylmethionine, phosphocreatine, and were negatively correlated with cholesterol and glycosphingolipid hexosylceramide (HexCer (d18:1/24:1); **Fig. 5B**). The most significantly changed analytes in the comparison of high vs. low mKate2 disease displayed on the heatmap (**Fig 5C**) included many of the energetic metabolites identified in the PC contributor analysis. These metabolites were significantly changed with consistent directionality driven by TREM2 levels in both healthy and disease mice shown in volcano plots (**Fig 5D**, **Extended Data Table 2**). Consistent with this pattern, free cholesterol (∼1.2 log_2_ FC) and palmitoylcarnitine (∼1.6 log_2_ FC) levels were both significantly (p<0.0005) lower in high vs. low TREM2 microglia, suggesting increased lipid catabolism and cholesterol efflux. To interrogate the concordance of the metabolites with TREM2 expression and disease status, we compared fold-changes in disease vs. healthy mice (**Fig 5E**). Many metabolites associated with positive cellular energetics such as s-adenosylmethionine (SAM), coenzyme Q, and creatine were increased in subpopulations expressing higher TREM2 level, in both a healthy and disease context. Visualization of the data along the axis of TREM2 expression per mouse shows that increasing TREM2 correlates with higher abundance of fructose-1,6-bisphosphate, sedoheptulose-7-phosphate, creatine, glutathione (GSH) and SAM, and reduced abundance of lactate. Taken together, this data suggests increased glucose uptake correlates with higher pentose phosphate pathway (PPP)^48^, methylation and redox capacity in high disease or mid healthy vs. low TREM2 expressing microglia (**Fig. 5F**). Overall, increased TREM2 levels are associated with elevated energetic and metabolic capacity, and improved cholesterol handling in microglia.

**Figure 5.**
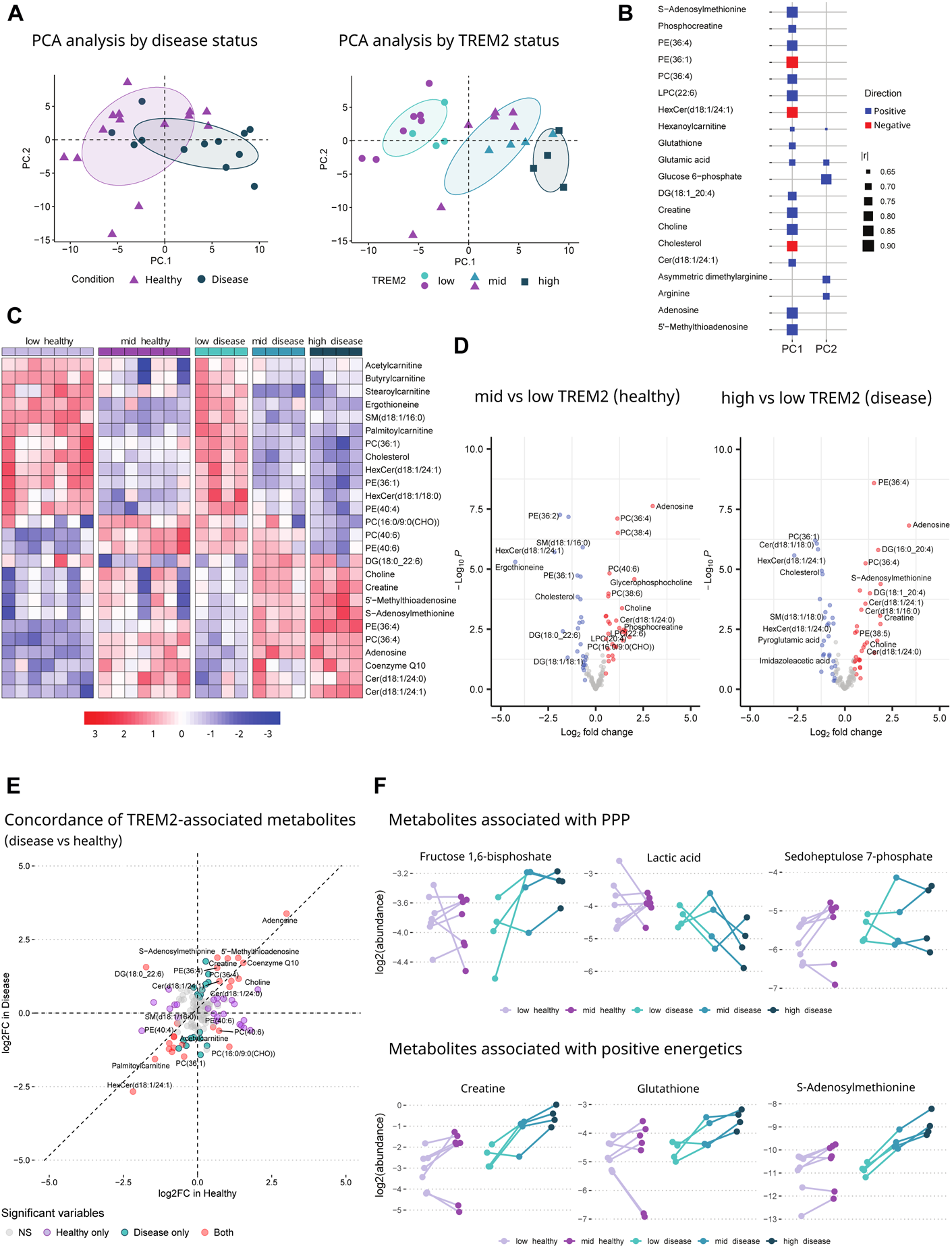
Microglial TREM2 levels correlate with cellular energetics metabolites and PPP intermediates. (A) PCA analysis of all 141 analytes measured in microglia sorted by TREM2 levels in healthy and disease mice, n = 4-7 mice per genotype, age = 9 months. (B) Top 20 highest contributors to PC1 and PC2 (lrl: correlation coefficient). (C) Heatmap displaying most significantly changed lipids/metabolites within the comparison of high vs. low TREM2 in disease mice. (D) Volcano plots showing significantly changed (p < 0.05 and log2(fold change) > 0.5) lipids/metabolites in sorted microglia by TREM2 levels in healthy and disease mice. (E) Concordance analysis of metabolites associated with TREM2 level differences in healthy vs. disease states. Significantly (p < 0.05 and log2(fold change) > 0.5) changed analytes are highlighted according to disease status, and those modified in both healthy and disease mice are labelled in red. (F) Abundances of metabolites associated with PPP and positive energetics vs. TREM2 levels plotted on a per mouse basis.

### A TREM2 agonist antibody induces metabolic changes dependent upon TREM2 expression level

We previously demonstrated that a mouse TREM2 agonist antibody (4D9)^34^ engineered to cross the blood brain barrier using an antibody transport vehicle (ATV:4D9) boosts TREM2 activity in vivo and increases microglial metabolism^25^. To determine the effect of this antibody on microglial subpopulations stratified by TREM2 expression levels, mice were dosed monthly with 1 mg/kg ATV:4D9 or isotype control for 4 months (**Fig. 6A, Extended Data Fig. 4A**). Here, *Trem2-mKate2^KI/wt^.APP^SAA/SAA^.hTfR^KI/KI^*mice were used to allow the antibody to cross the blood brain barrier by TfR binding^25,49^, as well as to validate our results in a second amyloid model. Mice received the last antibody dose 1 month prior take down. In addition, 30 mg/kg of fluorodeoxyglucose (FDG) was dosed 1 hour before animal takedown to enable detection of phospho-FDG (pFDG) in sorted microglia (see **Extended Data Fig. 4B** for validation of the method). After perfusion, brains were dissociated, microglia isolated and sorted by flow cytometry into low, mid, and high TREM2 expressing subpopulations (**Extended Data Fig. 4C**); and cell extracts were then subjected to LC-MS (**Fig. 6A**). PCA analysis and loading scores (**Fig. 6B, 6C**) of the LC-MS data show that the low TREM2 subpopulations derived from mice dosed chronically with ATV:4D9 and isotype control have similar lipodomic and metabolomic profiles, while the mid and high TREM2 subpopulations are distinct and cluster depending upon antibody dosing. Interestingly, high TREM2 isotype and mid TREM2 ATV:4D9 cluster together. Likewise, mid TREM2 isotype and high TREM2 ATV:4D9 form one cluster (**Fig. 6B**), suggesting that the TREM2 agonist antibody differentially modulates microglial metabolism dependent on TREM2 expression level. The component separation is driven by phospholipids such as PCs and PEs, as well as glycolysis intermediate pFDG and lysosomal BMP lipid species. Consistent with PCA analysis, the heatmap (**Fig. 6D**) and volcano plots (**Fig. 6E, Extended Data Table 3**) suggest that upon chronic dosing of ATV:4D9, very few significant changes are induced in the low TREM2 subpopulation, and the metabolic changes induced by ATV:4D9 are predominantly observed in the mid and high TREM2 expressing microglia (**Fig. 6D, 6E, Extended Data Table 3**). In contrast to the low TREM2 expressing microglia, mid TREM2 levels were associated with significant increases in key metabolic indicators of glycolysis (pFDG), lysosomal function (BMP(22:6/22:6)) and peroxisomal function (PE-(P16:0/20:4)) (**Fig. 6F**). Furthermore, ATV:4D9 treatment was highly effective in increasing levels of polyunsaturated fatty acid (PUFA)-containing phospholipids, including sub-species of phosphatidylcholine (PC), phosphatidylethanolamine (PE), and phosphatidylserine (PS) in the mid-TREM2 subpopulation (**Fig. 6F, Extended Data 5D**). Interestingly, while in the mid-TREM2 expressing microglia ATV:4D9 mediated increased levels of lipids and metabolites, the trend was reversed in the high-TREM2 expressing microglia (**Fig. 6D-6F**), supporting the above PCA analysis (**Fig. 6B**). These findings indicate that the level of TREM2 expression on the cell surface of microglia is critical for mediating the metabolic response to the agonist antibody.

**Figure 6.**
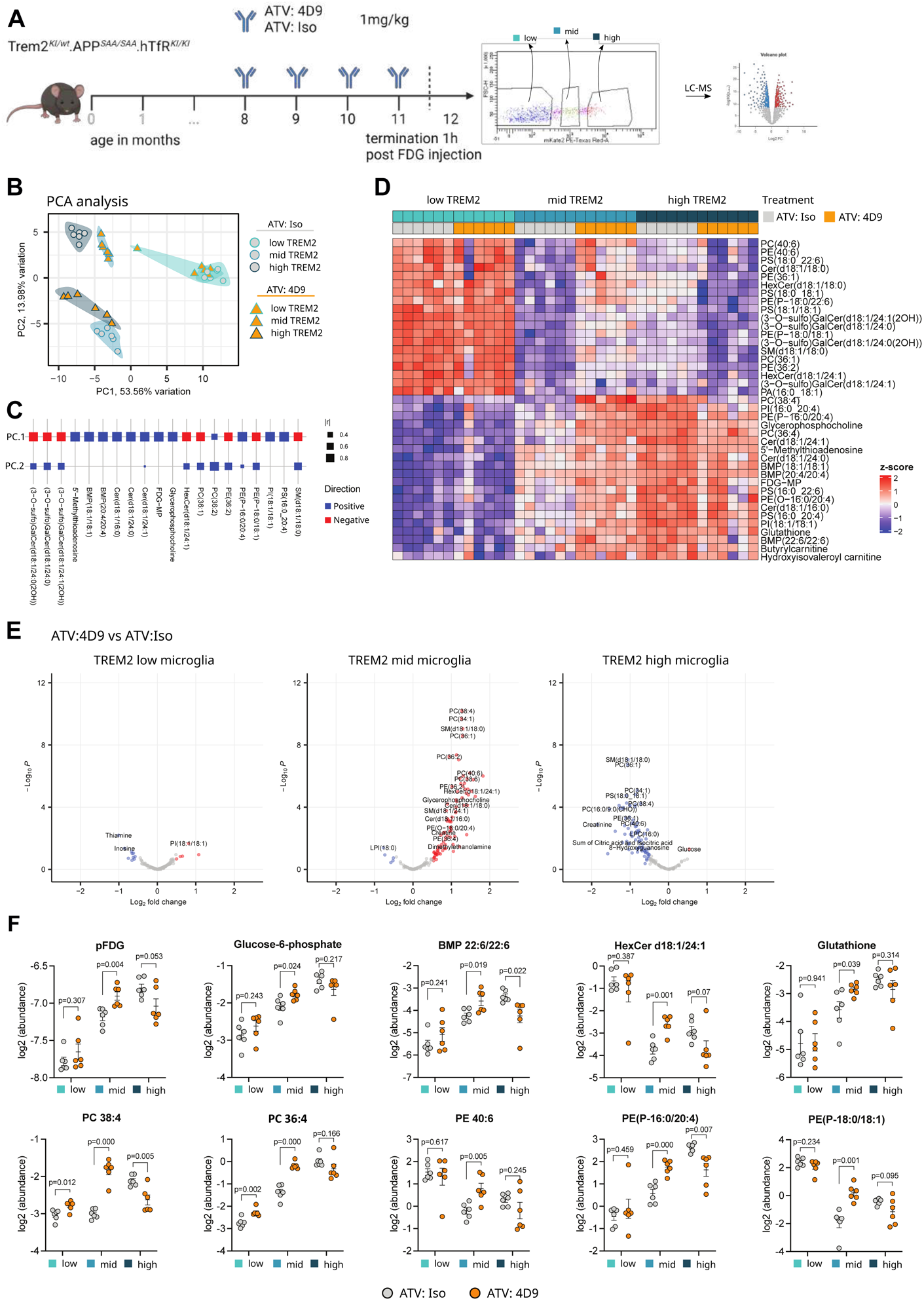
TREM2 agonist ATV:4D9 modulates metabolic profile dependent upon TREM2 expression level. (A) Schematic of experimental design. 8-month-old *Trem2-mKate2^KI/wt^.APP^SAA/SAA^.hTfR^KI/KI^*mice were injected with 1 mg/kg ATV antibody monthly until 11 months. At 12 months, mice were injected with 30 mg/kg FDG, perfused 1 hour later and then sorted into low, mid and high mKate2 expressing populations. Cells were pelleted and frozen for further analysis n = 6 mice per genotype, age = 12 months (B) PCA analysis of all 149 analytes measured by LC-MS in sorted microglia. (C) Top 20 highest contributors to PC1 and PC2 (lrl: correlation coefficient). (D) Heatmap of top 40 differentially regulated lipids and metabolites. (E) Volcano plots of treatment comparisons within the same TREM2 level. (F) Feature plots of individual lipids/metabolites altered by ATV:4D9 (orange circles) vs ATV:Iso (grey circles) within TREM2 subpopulations (low, mid, high) (Mean ± SEM, p-value calculated within each group by unpaired Student’s t-test).

## Discussion

Our study supports the idea that microglia can readily adjust their energy metabolism to resolve challenges^46^. We have shown previously that microglia have a high rate of glucose uptake and that this is dependent on the activation status of the cells^24^. Using single cell radiotracing, we demonstrate that microglial glucose uptake is strongly correlated with the expression level of TREM2. By combining glucose uptake studies with metabolomics, we also ascertain the likely fate of glucose in supporting the cellular redox environment required for glutathione and lipid synthesis. Microglia express the hexokinase 2 (HK2) isoform which preferentially diverts glucose for NADPH generation required for glutathione redox and lipid biosynthesis via the pentose monophosphate pathway (PPP)^50,51^. Although NADPH was not directly measured due to technical challenges, this interpretation is supported by our observations that 1) increased PPP intermediates occur at the expense of lower lactate, 2) higher abundance of several unsaturated long chain PE and PC species, 3) increased total glutathione levels and 4) TREM2-associated increase in transcripts of lipid biosynthesis enzymes. In addition, cellular S-adenosylmethionine (SAM) which is required for PC synthesis^52^ was significantly upregulated in high vs low TREM2 expressing microglial subpopulations. Increased glutathione is inversely associated with pro-inflammatory potential by lowering ROS production^53^, supporting cellular proliferation^54^, and decreasing ferroptosis potential^55^ in microglia. Furthermore, treating mice with the TREM2 agonist ATV:4D9 resulted in metabolic alterations akin to those observed along the axis of increasing endogenous TREM2 levels particularly in mid-expressing microglia. Interestingly, in high TREM2 expressing microglia ATV:4D9 resulted in attenuated metabolic profiles like those observed in mid-TREM2 expressing microglia with isotype treatment. These results demonstrate that TREM2 agonism can regulate microglial metabolic activity, while tempering activity in high-TREM2 expressing microglia. These findings suggest that TREM2 levels are a key regulator of metabolic state and further demonstrate that TREM2 has an immunomodulatory role whereby it can tune metabolic activity up or down depending upon a threshold of the TREM2 receptor. Increased accumulation of PUFA, PC, PE and plasmalogens suggests that ATV:4D9 could improve cell membrane flexibility that is required to support higher phagocytic and lipid clearance activities. Notably, plasmalogens, an abundant class of phospholipids, have been found to be deficient in AD patient brains, and some studies have supported beneficial effects of reduced inflammation and neuroprotection with plasmalogen administration^56–58^.

Taken together, TREM2-associated increases in glucose and its usage to support glutathione redox may be a key underlying mechanism by which TREM2 promotes microglial survival in times of stress. Our lipidomic and metabolomic data strongly suggest that TREM2 expression is a driver of lipid catabolic pathways, including fatty acid oxidation (FAO), which is consistent with the previously reported ability of TREM2 agonist antibodies to clear lipid droplets and promote FAO in iPSC-derived human microglia^25^. Importantly, largely based on KO studies, TREM2 has been identified as a major regulator of microglial cholesterol transport and metabolism^28,29,59^. Our study lends strong support to this notion, by showing lower free cholesterol levels in TREM2 high microglia, consistent with higher catabolism or efflux capacity. Since cholesterol storage forms, like cholesteryl esters, are known to accumulate in AD brain^60^, the impact of the TREM2 pathway on cholesterol clearance is likely to be disease-relevant. Since TREM2 risk variants are thought to be loss-of-function, TREM2-driven glucose uptake is likely perturbed in individuals carrying these variants, which is supported by previous studies of the T66M mutation in mice^23^. This is in line with reduced cerebral glucose metabolism and regional hypometabolism that has been reported in an Nasu-Hakola disease patient^61^. There is a high prevalence of AD and metabolic disease being co-morbidities, as well as risk factors for each other ^62–64^, hinting towards the disease relevance of microglia metabolism. Moreover, glucose hypometabolism may be a key manifestation of prodromal AD and may therefore even be useful as a biomarker. However, direct comparison of these results should be conducted with caution, as the functionality of macrophages in the periphery and microglia in the brain can be diverse given the different environments. Surprisingly, we only detected minor differences in disease vs. healthy conditions, while the TREM2 level effects were largely concordant between the conditions. These findings indicate that regardless of brain environment, TREM2 can function as a metabolic driver to increase cellular bioenergetic capacity, likely in accordance with increased demands for innate immune functions.

Subjecting the TREM2 stratified subpopulations to bulk RNAseq, we validated that sorting based on mKate2 levels is indeed a proxy for TREM2 expression differences. Using the published PIG signature by Chen et al ^42^, we performed a gene set variation analysis assessing the data in a spatial context. In contrast to Chen et al ^42^, isolated microglia displayed a significant upregulation of the PIG signature with increasing TREM2 levels. This effect is enhanced in the disease context due to plaque presence, however, we also observed a decreased enrichment score comparing mid healthy to low healthy microglia. This suggests that the PIG signature reflects a general microglial response to an environment requiring higher phagocytic capacity and not necessarily an amyloid-specific microglial response. While plaque vicinity constitutes an environment in which the signature has reached its maximum, it is dependent upon TREM2 upregulation. Hence, we postulate that the PIG signature reflects a cellular response enabling subpopulations of microglia to perform activities such as removing myelin to maintain brain homeostasis^65^. Further comparing the gene signatures of the subpopulations to homeostatic, stage 1 DAM, and stage 2 DAM as defined by Keren-Shaul et al^7^, we found that the majority of microglia in the healthy mice expressed a homeostatic signature together with the minor presence of a DAM 1 signature which can be potentially explained by aging-related myelin damage that microglia clean-up ^23,66,67^. Strikingly, an increasing fraction of the DAMs, specifically of stage 2 DAMs, was observed in disease microglia. This finding aligns well with the enrichment of the PIG signature in the disease context pointing towards higher plaque proximity required for the emergence of DAMs ^7^ dependent on the *Trem2* level but also suggests that the expansion of the *Trem2*-dependent stage 2 DAM population could be additionally regulated by the *Trem2* level. Additionally, we validated the reporter-based sorting approach by comparing the signatures to the classifications from Grubman et al^43^. Here, we saw a highly consistent pattern in which the disease conditions showed the expected upregulation of the 5xFAD Methoxy-X04 positive signature with a concomitant WT signature reduction. To gain further insights into the functional state of the microglia with different TREM2 levels in the health and disease condition, gene co-expression network analysis was performed identifying six gene modules. One of the modules, plum, contained type I and II IFN response genes, which were specifically elevated in the disease context and in particular in high TREM2 microglia. A previous study recently identified a subset of DAM expressing increased levels of type I IFNs promoting cellular states detrimental to memory and cognition ^68^, but the upregulation of type I IFN in disease microglia based on plaque proximity and *Trem2*-level has not been documented, yet ^68^. Additionally, type II IFN is known to prime microglia for the induction of enhanced secretion of inflammatory molecules such as IL-6 and TNFα and giving rise to neurotoxic phenotypes that drive neurodegeneration ^69^. Future studies are needed to understand the role that TREM2 levels and type I and II IFN signalling play in disease progression including studies addressing spatial distribution of this microglial subpopulation.

Previous studies suggested homeostatic genes such as *Cx3cr1* and *P2ry12* are suppressed in disease-associated microglia^7,8^, as well as following phagocytosis of apoptotic neurons by microglia^8^. Here, we link this information to TREM2 levels as we demonstrate the homeostatic signature to be gradually downregulated in disease with increasing TREM2 expression. Proximity to amyloid plaques, an increased expression of stage 2 DAM genes, and/or the high expression of type I and II IFNs can cause a high energetic demand on microglia. Supporting our lipidomic analysis, we observed a TREM2-dependent upregulation of genes associated with oxidative phosphorylation and lipid metabolism (e.g. the lightgreen and steelblue modules, Fig. 3G-I). Overall, these findings extend the current understanding of TREM2-dependent regulation of microglial state and metabolism with a higher resolution regarding the TREM2 expression level. We show that physiological TREM2 expression and function occurs as a gradient as opposed to an on/off switch.

By treating *Trem2-mKate2^KI/wt^.APP^SAA/SAA^.hTfR^KI/KI^* mice chronically with the brain penetrating version of the well characterized TREM2 agonistic antibody ATV:4D9^25,34^ we could demonstrate that microglia with intermediate expression levels of TREM2 reacted the most by increasing metabolites related to glucose uptake as well as antioxidants, reflective of an overall increased metabolic fitness. Some metabolites decreased in high vs mid subpopulations upon chronic dosing, suggestive of a ceiling effect. Similarly, too little expression of TREM2 may not provide enough target for the TREM2 agonist to be effective, further demonstrating the importance of the TREM2 expression level. These findings may have important consequences for the current design of clinical trials using TREM2 agonists. Based on our findings, one must carefully monitor TREM2 expression in patients, to determine the optimal time point for interference. Whether soluble TREM2 in the cerebrospinal fluid^70^ can be used as a biomarker, needs to be determined.

Taken together, our data highlights the correlation of TREM2 expression with microglial transcriptome and metabolic activity. Further, we provide evidence that transcriptional programs linked to differential levels of TREM2 are further shaped by the disease state, here exemplified in an amyloid plaque disease model. Importantly, we found that TREM2 surface expression is the major driver for microglial metabolic profiles, which could be modulated further by a TREM2 agonistic antibody. Given that we observed TREM2 upregulation to be gradual in plaque proximity as well as the bidirectional modulation by the TREM2 agonistic antibody, it is important to determine the optimal timepoint for therapeutic intervention in patients.

## Methods

### Expression vector design and expression

To create the reporter mouse, we developed a plasmid-based construct. mKate2 was chosen as a fluorescent tag^35^. To ensure that the tag does not interfere with normal TREM2 expression, function or localization we inserted the P2A peptide sequence, which allows the separate expression of both proteins^36^. Additionally, we included a KDEL sequence after the mKate2 sequence to retain it in the endoplasmic reticulum (ER)^37^. To create the expression vector, we obtained genomic mouse DNA from C57Bl6/J animals from which the TREM2 sequence was amplified. We obtained the mKate2 plasmid from Addgene and amplified the sequence by standard polymerase chain reaction (PCR). Primers for amplification were designed with overhangs compatible with the Gibson assembly cloning. PCR products were purified by gel extraction before being cloned into a pcDNA3.1 vector.

### Genetic editing of mice

All animal experiments were approved by the Ethical Review Board of the Government of Upper Bavaria. Mice were group housed with littermates on a normal 12 hour light/dark cycle with *ad libitum* access to food and water. Both genders were used for all experiments. The mKate2 cassette was targeted to the area between exon 5 and the stop codon of the endogenous murine *Trem2* gene using a CRISPR/Cas9 strategy (**Fig. 1C**). Pronuclear microinjections were performed to integrate the desired reporter sequence into C57Bl/6J mouse zygotes. Offspring was screened for the reporter cassette by PCR. Sequencing was performed on offspring with successful knock in to ensure there were no mutations in the insertion. Additionally, the most likely off target loci were sequenced before selecting the optimal animals to establish the colony. Homozygous *Trem2-mKate ^KI/KI^*mice were crossed with *APP/PS1 mice* ^41^ to generate transgenic *Trem2-mKate2^KI/wt^;APP/PS1* (hereafter referred to as disease mice) as well as non-transgenic *Trem2-mKate2^KI/wt^*mice (hereafter referred to as healthy mice). For the experiments in which we dosed with the agonistic antibody, the reporter mice were crossed to *App*^SAA^ knock in^46^ as well as *hTfR* knock in (human transferrin receptor expressing) mice. The experimental mice expressed the reporter heterozygous while *App*^SAA^ and the *hTfR* knock in were homozygous (*Trem2-mKate2^KI/wt^.APP^SAA/SAA^.hTfR^KI/KI^*).

### Traumatic brain injury

Controlled cortical impact (CCI), a model of brain injury was performed as described previously^38,71^. Briefly, *Trem2-mKate2* reporter mice were injected with buprenorphine (0.1 mg/kg) before being anesthetized with isoflurane (4%, 3 seconds). The CCI was achieved by using a pressure-controlled impactor (L. Kopacz, University of Mainz) with the following settings: impact velocity 6.5m/s, 0.5 mm penetration depth and 150 ms contact time. The injury was delivered directly to the dura after a right parietal craniotomy had been performed. The exposed dura was sealed using tissue glue (VetBond™, 3M Animal Care Products). During the whole procedure, anaesthesia was maintained with 1,5-2,5% isoflurane in oxygen/air and the animals heart rate and body temperature was monitored continuously. After the surgery, animals were allowed to recover in a heating chamber at 34°C with 60% humidity, to avoid hypothermia. Mice were monitored, scored and injected with carprofen (5 mg/kg) for 3 days following the surgery. 72 hours post impact animals were sacrificed and perfused, alongside naïve animals which had not undergone the procedure.

### Perfusion

At indicated ages mice were anesthetized by injection of Ketamine/Xylazine mix or CO_2_ inhalation before being transcardially perfused with ice-cold PBS. The brain was removed and separated into hemispheres. One half was snap frozen for biochemical analysis and the other half was post fixed by immersion in cold 4% paraformaldehyde (PFA) for histological assessment. For FACS analysis, mice were perfused with PBS, olfactory bulb and cerebellum were removed and the remaining brain was used to isolate microglia. At indicated times, mice received a single dose of 30 mg/kg FDG (Sigma) 1 hour prior take down.

### Immunofluorescence

Mouse tissue was changed from 4% PFA to PBS the day after perfusion. 50 µm thick sagittal brain sections were cut on a vibratome, collected in PBS and stored at 4°C until needed. Sections were transferred to 24-well plates and incubated with a blocking buffer (BB) containing 0.5% BSA and 0.5% TritonX-100 in 1x PBS, for 1 hour at room temperature (RT) with slow agitation. Primary antibodies (**Extended Data Table 4**) were diluted in BB and incubated overnight at 4°C again with slow agitation. The next day sections were washed 3x 10 minutes in 1x PBS before Alexa conjugated secondary antibodies (1:500) and DAPI (1:1000) (**Extended Data Table 4**), diluted in BB were added. Sections were incubating in the dark for 1 hour at RT while shaking. Sections were washed again 3x 10 minutes before being mounted on glass slides and cover slipped using Fluoromount-G (Invitrogen). Slides were sealed with clear nail polish and allowed to dry for at least 24 hours in the dark before being imaged.

HeLa cells were grown on cover slips, transiently transfected with lipofectamine 2000 (Invitrogen) for 48 hours and washed 3x with PBS before being fixed with 4% PFA for 20 minutes at RT. Cells were permeabilized for 10 minutes at RT with PBS containing 0.2% Triton-X. Cells were washed in PBS 3x 10 minutes and then blocked in 5% serum for 1 hour at 37°C. Primary antibody) (**Extended Data Table 4**) was diluted in BB and incubated at RT for 2 hours. After washing, cells were incubated with secondary antibodies and DAPI for 1 hour at RT. Cells were washed once more before being mounted on glass slides. Images were taken with a LSM800 (Zeiss) Confocal microscope. Laser power and gain were kept consistent across imaging sessions. The Zen blue software (Zeiss) or ImageJ were used for quantification.

### qPCR

RNA isolation was performed using the RNeasy Mini kit (Qiagen), following the manufacturer’s instructions. Briefly, either harvested cells or mouse brain powder was resuspended in RLT buffer (Qiagen) for cell lysis and transferred onto spin columns. Optimal DNase I treatment was performed for 15 minutes at RT. RNase-free water was used to elute the RNA and concentrations measured using a Nanodrop. RNA was kept at -80°C or directly used for cDNA generation using the Superscript IV reverse transcriptase protocol (Invitrogen). TaqMan technology (Applied Biosystems) was used to analyse mRNA expression. The 7500 Fast Real-Time PCR system (Applied Biosystems) was used.

### Microglia isolation for FACS

Microglia isolation was performed using the MACS system (Miltenyi Biotec) as described previously ^72^. However, no CD11b enrichment step was performed as the intrinsic mKate2 reporter is only expressed in TREM2 expressing (microglia) cells. Hence, FACS was performed based on the mKate2 fluorescence signal and sorted into low, mid and high expressing subpopulations, as a proxy for the TREM2 expression of individual cells. The sorted fractions were spun down and the pellets stored at -80°C before being processed for further analysis, as described below.

### Transcriptomic analysis

For a subpopulation of microglia RNA sequencing was performed. To this purpose, cells were sorted as described above and immediately stored in Trizol. RNA extraction was performed as previously described. For RNA sequencing, both the RNA quantity and integrity were assessed using the HS RNA assay on a Tapestation 4200 system (Agilent). The Smart-seq2 protocol, as described by Picelli et al. (2014)^73^ was used for the generation of non-strand-specific, full transcript sequencing libraries. Briefly, 1 ng of total RNA was transferred to buffer containing 0.2% TritonX-100, protein-based RNase inhibitor, dNTPs, and oligo-dT oligonucleotides to prime the subsequent RT reaction on polyadenylated mRNA sequences. The SMART RT reaction was conducted at 42°C for 90 minutes using commercial SuperScript II (Invitrogen) and a Template-Switching Oligo. A pre-amplification PCR of 16 cycles was carried out to generate double-stranded DNA from the cDNA template. Subsequentely, 100 pg of amplified cDNA were used for tagmentation and enrichment using the Nextera XT kit (Illumina) to construct the final sequencing libraries. Libraries were quantified using the Qubit HS dsDNA assay, and library fragment size distribution was determined using the D1000 assay on a Tapestation 4200 system (Agilent). The sequencing was performed in single-end mode (75 cycles) on a NextSeq 500 System (Illumina) with NextSeq 500/550 High Output Kit v2.5 (150 Cycles) chemistry. The sequenced reads were aligned against the Gencode mouse reference genome vM16 using kallisto (0.44.2)^74^ and samples with more than 50,000,000 reads were downsampled to 50 % of the original number of reads. The following analysis steps were performed in R (v4.1.0)^75^ and R Studio (v1.4.1717)^76^. The count matrix was imported into the analysis pipeline using R/tximport (v1.20.0)^77^. Genes with less counts than number of samples were excluded from the analysis resulting in 26,492 genes kept in the analysis. Normalization of the count matrix was computed with R/DESeq2 (v1.32.0)^78^ and a variance stabilizing transformation applied using the DESeq2 rlog function at default settings. Differential expression analysis based on the DESeq2 package was performed adjusting p-values according to independent hypothesis weighting from the IHW package (v1.20.0)^79^ and applying normal shrinkage. DEGs were defined based on a fold change threshold > 2 and a p-value threshold of < 0.05. Cell type abundances of the variance stabilized data were determined by CIBERSORTx (https://cibersortx.stanford.edu/)^80^ with default parameters using two microglia single-cell RNA-seq data sets (GSE98969 ^7^, GSE165306 ^43^) as reference. Signature enrichment was calculated on the variance-stabilized data per sample using R/GSVA (v1.40.1)^81^ with default settings and a Wilcoxon rank sum test with subsequent Benjamini-Hochberg adjustment was applied with R/rstatix (v0.7.0, https://CRAN.R-project.org/package=rstatix) to test for significance. Permutation tests were computed by drawing 500 random, unique signatures of the same size as the signature being assessed. Gene set variation analysis (GSVA) enrichment scores were calculated for each gene signature and an ANOVA followed by Benjamini-Hochberg adjustment was applied over the 5 microglia subpopulations. The likelihood of the enrichment result of the tested signature was computed by dividing the number of random gene signatures with lower adjusted p-values by the number of permutations. To assess the signature enrichment on a population level, expression level statistics from a non-parametric kernel estimation of the cumulative density function of each variance-stabilized gene expression profile per sample were calculated^81^ prior to computing the mean for the 5 different microglia subpopulations. A gene set enrichment analysis (GSEA) was performed with the transformed data as the input using R/fgsea (v1.18.0)^82^. The gene co-expression network analysis was performed using R/hCoCena (v1.0.1)^83,84^ on the normalized count matrix. Gene pairs with a Pearson’s correlation coefficients lower than 0.879 were excluded resulting in a network with 4,259 genes, 207,764 edges and an R^2^-value of 0.748. The Leiden clustering algorithm was used to identify 6 gene modules with a minimum size of 50 genes containing a total of 4,002 genes. Gene Ontology (GO)^85,86^ and the Molecular Signature Database (MSigDB) ^87,88^ Hallmark gene set enrichment was performed on the module genes making use of the functions included in the hCoCena workflow with default parameters. The bulkRNAseq data is available under the following GEO accession number: GSE271074

### Single cell radiotracing

Animals (15 months old) were food deprived for 2-3 hours before the experiment started. Mice received a single tail vein injection of 45 ± 3 MBq [^18^F]fluor-desoxyglucose (FDG)^24,47^. 30 minutes post injection animals were sacrificed by cervical dislocation and their brains removed, as described previously ^47^. A single cell suspension was generated, microglia were incubated with 10 µl of CD11b MicroBeads (Miltenyi Biotec) per 10^7^ total cells for 15 minutes at 4°C. Cells were washed, centrifuged at 300 g for 10 minutes, the pellet resuspended in 500 µl of D-PBS–0.5% BSA and then added to pre-wetted LS columns (Miltenyi Biotec). The columns were washed with 3 × 3 ml of D-PBS–0.5% BSA buffer after which the cells were flushed out using 5 ml of D-PBS–0.5% BSA buffer. The suspension was centrifuged and the remaining cell pellet resuspended in 1 ml of cold D-PBS and 10 µl DAPI staining solution. Using a MoFlo Astrios EQ cell sorter (B25982, Beckman Coulter), microglia were sorted based on their mKate2 expression into the subpopulations as described in the previous section. Sorted cell pellets were analysed in a gamma counter (Hidex AMG Automatic Gamma Counter) with decay correction to time of tracer injection for final activity calculations. The measured radioactivity (Bq) of the cell pellet was divided by the specific cell number in the pellet, to calculate radioactivity per single cell. Radioactivity per cell was then normalized by injected radioactivity and body weight^89^.

### Metabolomic and Lipidomic analyses

Metabolomic and lipidomic analyses were performed as described previously^46^, with some modifications. Sample extraction: Pellets from the sorted subpopulations were reconstituted on ice in 9:1 MeOH:water including internal standards, vortexed for 1 minute, and spun down for 5 minutes at 10,000 g. Supernatant was transferred to glass vial for analysis by LC-MS. Polar metabolites in electrospray positive mode: Targeted analysis of 275 polar metabolites was performed on an Agilent 1290 UPLC system (Agilent) coupled to Sciex triple quadrupole mass spectrometer (QTRAP 6500+, Sciex). All other settings were as described in Xia et al^46^. Polar metabolites in electrospray negative mode: Samples were dried and reconstituted in same volume of 9:1 MeOH:water fortified with 0.1% formic acid. Acidified samples were chromatographically resolved on an Imtact Intrada Organic Acid column (3 µm, 2x 150 mm, Imtakt) using a flow rate of 0.2 ml/minute at 60°C. Mobile phase A consisted of acetonitrile/water/formic acid = 10/90/0.1%. Mobile phase B consisted of acetonitrile/100 mM ammonium formate= 10/90%. The gradient was programmed as follows: 0.0–1.0 minutes at 0% B; 1.0–7.0 minutes to 100% B; 7.1 at 0% B; and 7.1-10 minutes at 0% B. Source settings used were as follows: curtain gas at 40 V; collision gas was set at medium [collision gas for TQ 6500+ settings was set at 8 psi]; ion spray voltage at -4500 V; temperature at 600°C; ion source Gas 1 at 50 psi; ion source Gas 2 at 60 psi; entrance potential at -10 V; and collision cell exit potential at -15.0 V. Lipidomics in positive and negative ionization modes: Targeted analysis of 153 lipids were performed as described in Xia et al^46^ but LC system used in this study was switched to the Agilent 1290 UPLC system (Agilent) with identical gradient settings. MS settings were unchanged. Data acquisition and analysis: Data acquisition was performed using Analyst 1.6.3 (Sciex) in multiple reaction monitoring mode (MRM) with optimized collision settings. Peak integration and quantification were performed using MultiQuant 3.02 (Sciex) software. Endogenous metabolites/lipids were expressed as area ratios to specific spiked in stable isotope internal standards. Peak annotation and integration were based on defined retention times and MRM properties of commercial reference standards. Of the 428 targeted analytes, 141 analytes passed QC, missing values, and outlier checks. Based on replicate analysis of pooled QC samples, mean coefficient of variance for across all analytes was 16.4%. For all analysis, raw area ratio data was log2 transformed and filtered to retain variables that are present in at least 70% of samples. Missing values were imputed using the kNN method using the 5 nearest donor variables. Unwanted variations in the dataset were adjusted for using the RUV4 method. Assessment of genotype (healthy vs disease) and TREM2 status (low, mid, high), a group factor was created by combining genotype and TREM2 expression level status. A linear mixed model was conducted with individual mice as a random effect, group status as independent factor, and RUV4 regression factors as covariates. Benjamini-Hochberg adjustment was used to adjust for multiple comparisons. Top contrast results were visualized using heatmaps, boxplots or volcano plots. All analyses were performed using R statistical software (version 4.0.2; R Core Team 2020).

### Statistics

Statistical analysis was carried out using the GraphPad Prism software. Student’s t-test (unpaired, two-tailed) was used to compare two groups, while one-way ANOVA (with Tukey correction) was used to compare multiple groups. A value of p < 0.05 was considered statistically significant. Statistical analysis of the Omics data was performed as described in the respective methods sections.

## Figures

***Extended Data Figure 1***

(A) – (C) qPCR expression levels of mRNA from *Trem2-mKate2* mice for total *Trem2*, *mKate2* and the *Trem2* wild type allele respectively; n = 2-5 mice per genotype. (D) Schematic representation of experimental groups of traumatic brain injury following controlled cortical impact. (E) – (H) qPCR expression levels of mRNA from *Trem2-mKate2* mice for total *Trem2, mKate2*, *Trem2* wild type allele, *Clec7a, Cd68* and *Grn* respectively; n = 2-6 mice per genotype.

***Extended Data Figure 2***

(A) Workflow of the bioinformatics analysis. (B) Gene set enrichment analysis (GSEA) of the PIG signature coloured by condition. Ranking of genes is based on expression level statistics and the running sum is visualized. (C) GSEA of an exemplary random signature coloured by condition. Ranking of genes is based on expression level statistics and the running sum is visualized. (D) Violin plot of gene set variation analysis (GSVA) enrichment scores of 500 random signatures coloured by condition. Statistics were computed by a Wilcoxon rank sum test followed by a Benjamini-Hochberg adjustment and a summary of all comparisons is displayed. (E) P-value distribution of GSVA enrichment results of 500 random, unique gene sets over all conditions on a log10 scale. Gene set size was based on the size of the PIG signature. P-values were computed with an ANOVA followed by a Benjamini-Hochberg adjustment. The red dashed line represents the adjusted p-value from GSVA enrichment of the PIG signature. (F) Stacked bar plot of microglial composition based on cell state deconvolution of the data set by Keren-Shaul et al. summarized per condition and coloured by activation state. Error bars represent the standard deviation. (G) Stacked bar plot of microglial composition based on cell state deconvolution of the data set by Grubman et al. summarized per condition and coloured by activation state.

***Extended Data Figure 3***

(A) Boxplot of the gene set variation analysis (GSVA) enrichment scores of AD-related and TREM2-related DEGs coloured by condition; n = 4-5 mice per condition, age = 9 months. Boxplots show the 25%, 50% (median) and 75% percentile, whiskers denote 1.5 times the interquartile range. (B) P-value distribution of GSVA enrichment results of 500 random, unique gene sets over all conditions on a log10 scale. Gene set size was based on the size of the AD-related and TREM2-related signatures, respectively. P-values were computed with an ANOVA followed by a Benjamini-Hochberg adjustment. The red dashed lines represent the adjusted p-value from GSVA enrichment of the AD-related and TREM2-related signatures, respectively. p-values calculated with Wilcoxon rank sum test with Benjamini-Hochberg adjustment.

***Extended Data Figure 4***

(A) Breeding scheme to generate the second amyloid mouse line *Trem2-mKate2.APP^SAA/SAA^.hTfR^KI/KI^*. Only one genotype was used and treated with either ATV: Isotype or ATV: 4D9 at 1 mg/kg; n = 6 mice per genotype, age treatment = from 8 until 11 months, sacrificed at 12 months. (B) Example graph of the level of pFDG that could be measured after dosing with different concentrations of FDG in microglia or astrocytes respectively. This graph is derived from method establishment with wildtype animals where no antibody treatment was performed. (C) Example contour FACS plots which were used to determine the gating strategy of antibody treated knock in animals to define and sort the microglia into low, mid and high. (D) Log fold changes in PUFA-containing phospholipids and plasmalogens by ATV:4D9 vs isotype in mid-TREM2 microglia.

## Declarations

### Ethics approval and consent to participate

All animal experiments were approved by the “Regierung von Oberbayern”.

### Availability of data and materials

All data analysed and interpreted in this study are included in this published article or its Extended Data files. The bulkRNAseq data is available under the following GEO accession number: GSE271074. The detailed LC-MS datasets generated during the current study are available from the corresponding author upon reasonable request.

### Author’s contributions

CH designed the study and applied for funding. AFF, KD, AR, BW, MBr, KMM, AC, JS and CH designed experiments. AFF, KD, AR, JHS, BvL, BW, LMB, KW, TU, ED, IK, BT performed experiments. WW, NP, MBr, MBe, GDP, JWL KMM, AC, JS and CH supervised the study. AFF, KD, AR, JHS, BvL, LMB, MBr, GDP, JWL, KMM, AC, JS and CH analysed and interpreted data. AFF, KD, JHS, BvL and CH wrote the manuscript. TU, KS, JJN, GDP, JWL KMM, and JS edited the manuscript. All authors read and approved the final version.

## Acknowledgments

The authors thank the animal facility staff and management of the Center for Stroke and Dementia Research for excellent animal care. Sonnet Davis and Jamal Alkabsh for their technical assistance with mass spectrometry data generation. We thank Pardis Khosravani and Benjamin Tast as well as the Core Facility Flow Cytometry at the Biomedical Center, Ludwig-Maximillians Universität München and the Core Facility Flow Cytometry at the University Hospital, Ludwig-Maximillians Universität München, for technical assistance with cell sorting.

