## Extended Data Figures and Tables for "TREM2 expression level is critical for microglial state, metabolic capacity and efficacy of TREM2 agonism"

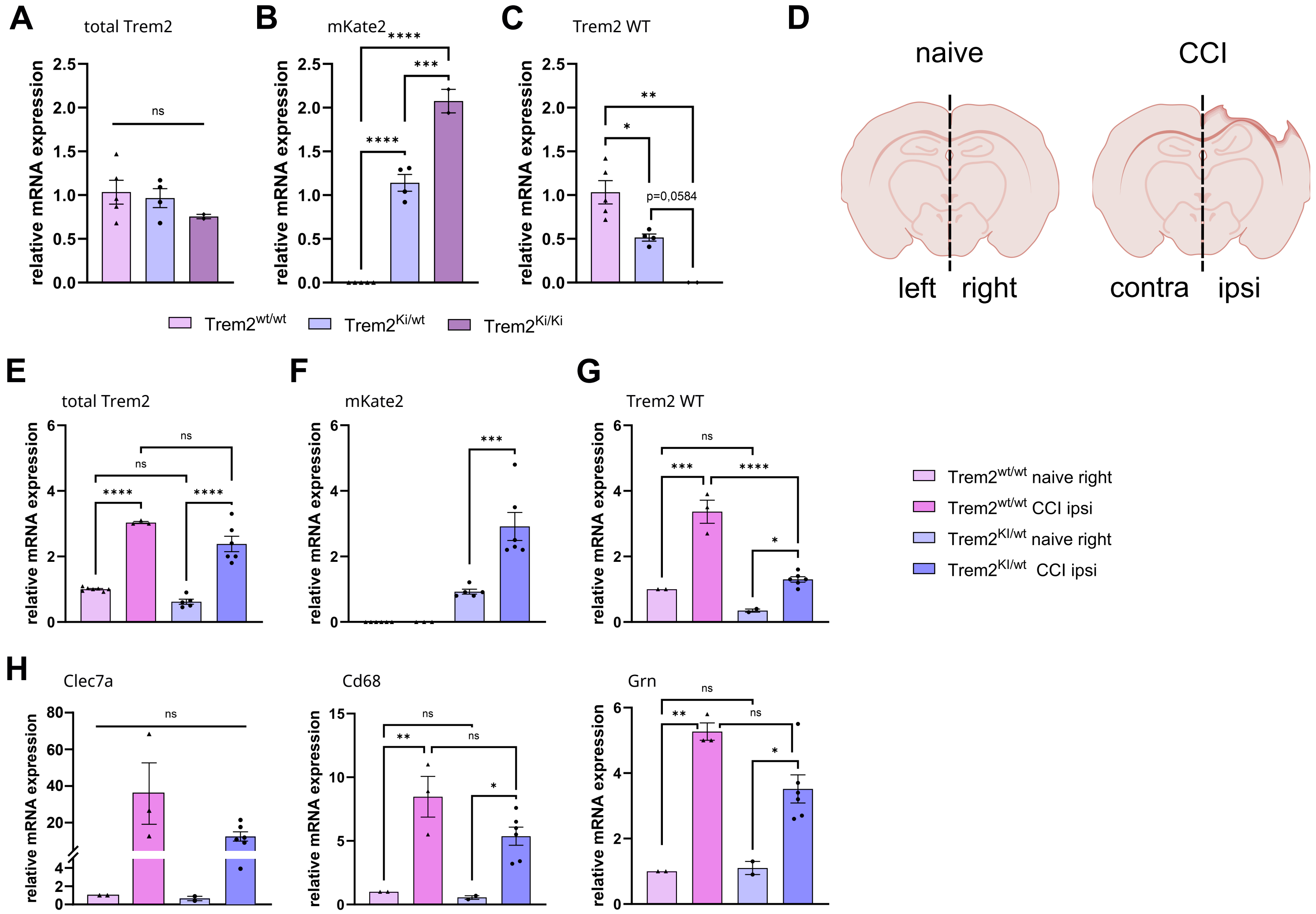

Extended Data Figure 1

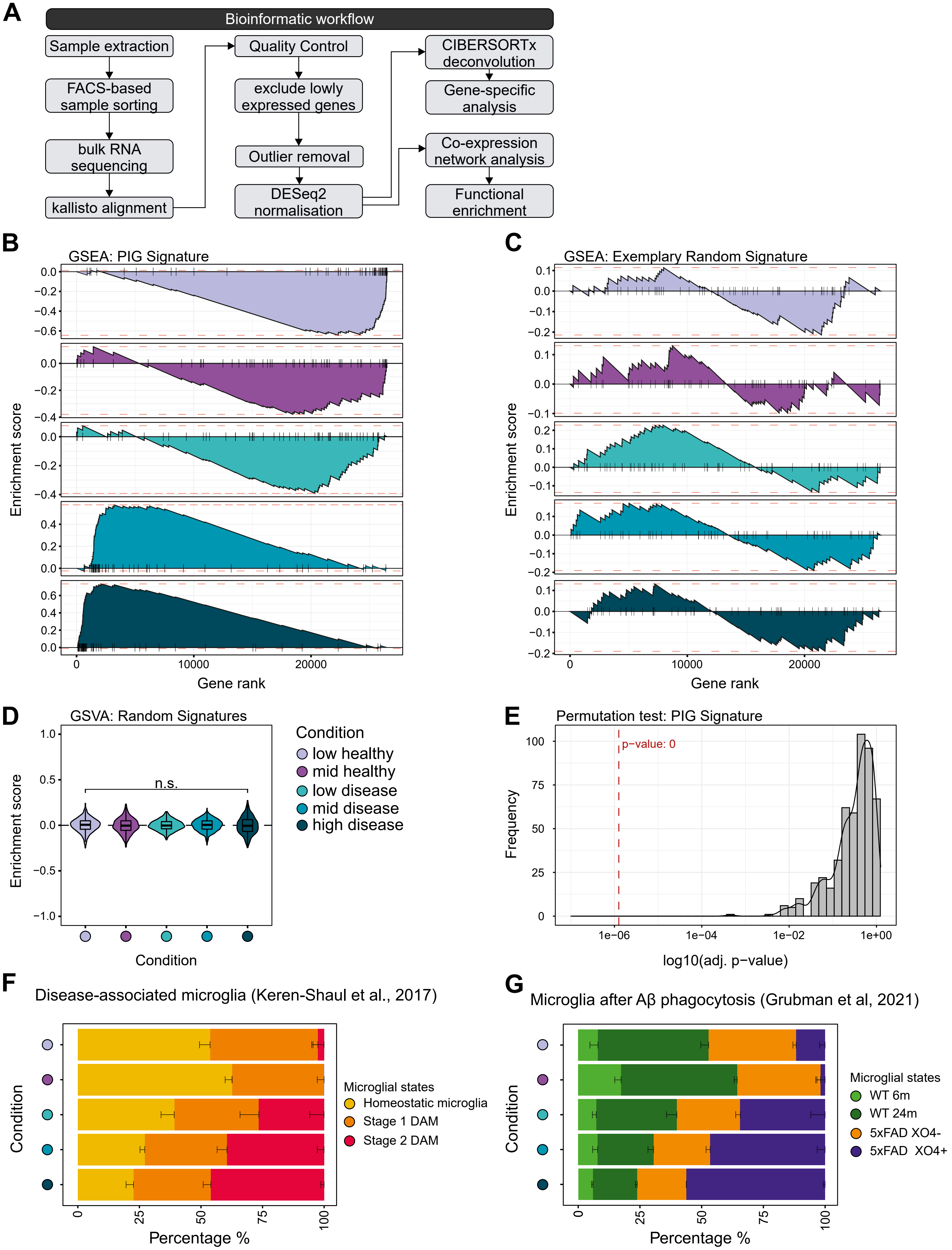

Extended Data Figure 2

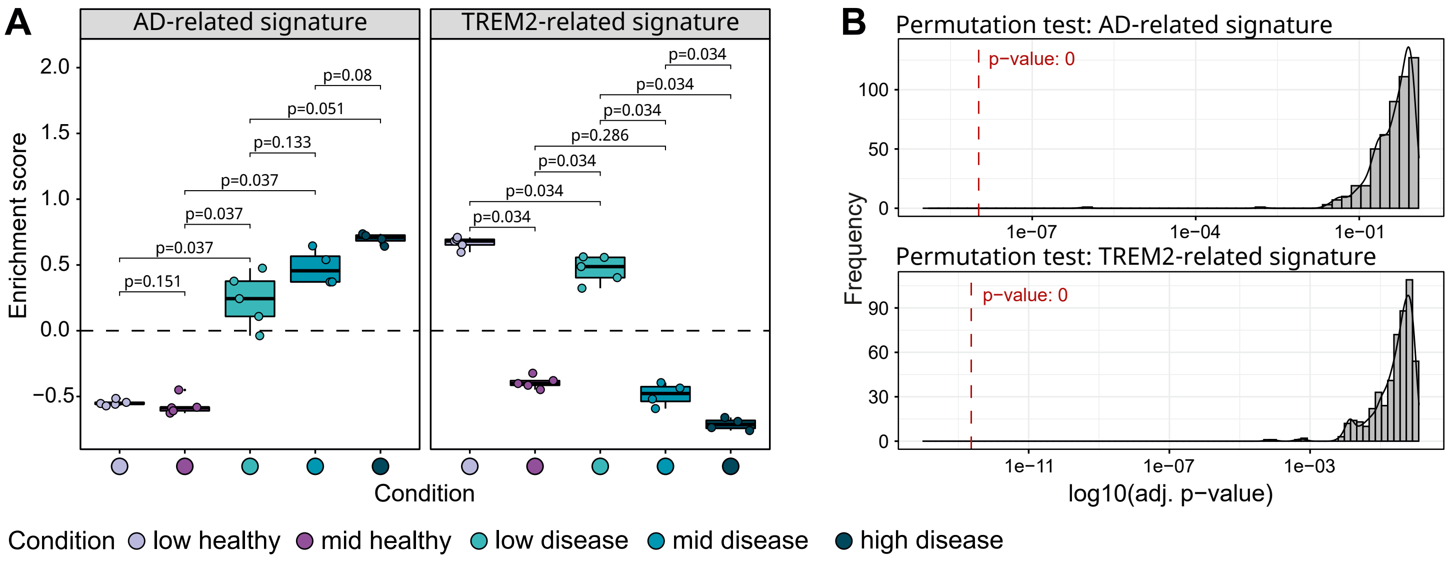

Extended Data Figure 3

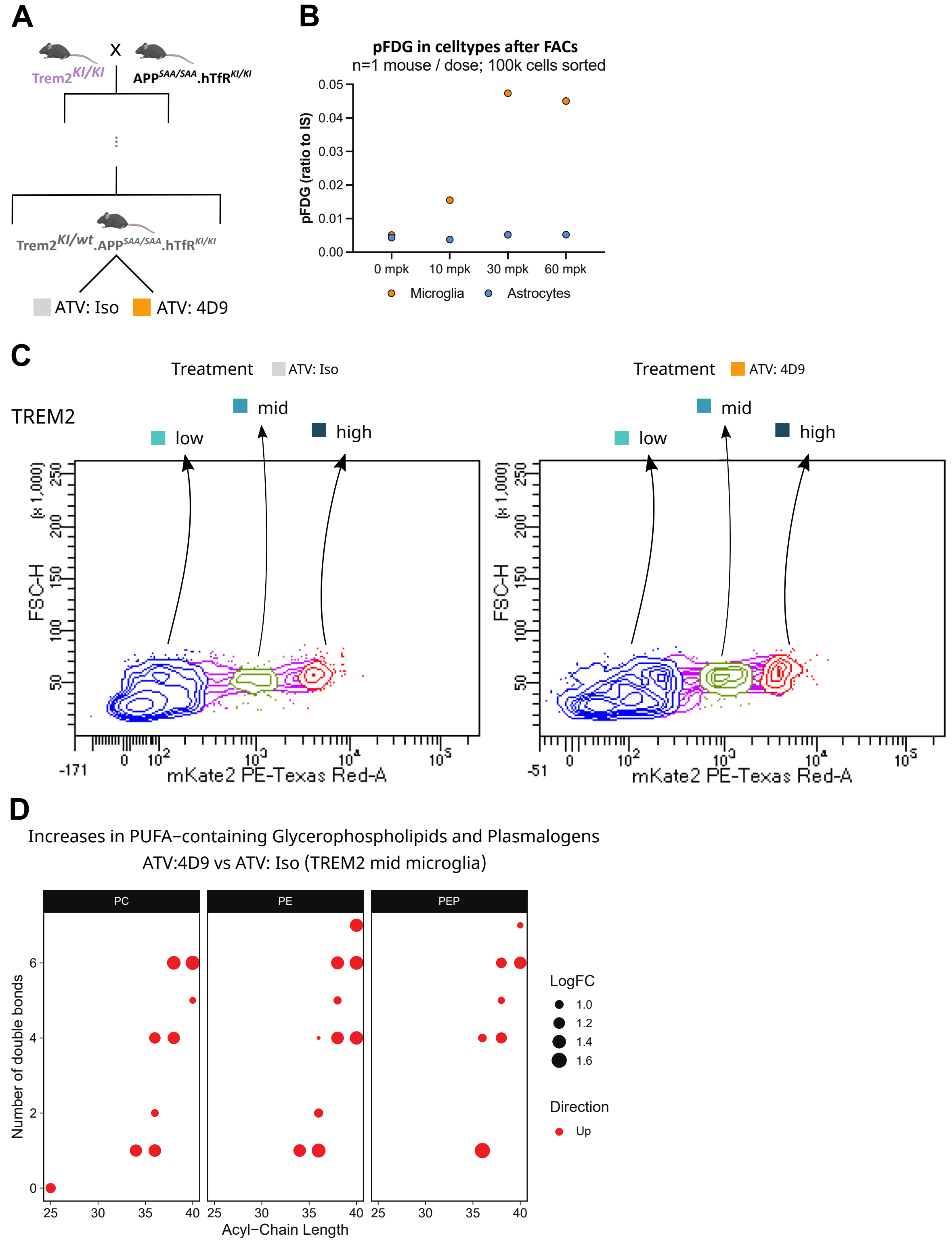

Extended Data Figure 4

| AD-related genes | Trem2-related genes |
|---|---|
| Pld3 | Clec10a |
| Ly9 | Cfp |
| Tcap | Napsa |
| Csf1 | Cd36 |
| Lpl | Slc1a2 |
| Rxrg | Eps8 |
| Cd34 | Mmp9 |
| Baiap2l2 | Mapt |
| Cdk1 | Ahr |
| Igf1 | Vnn3 |
| Madcam1 | Ntsr2 |
| Efr3b | Aoah |
| Gfap | Iqgap2 |
| Hif1a | Apod |
| Rai14 | Dtna |
| Ank | Ltb |
| Dpys | Vldlr |
| Tmem178 | Slco3a1 |
| Lox | Rgs18 |
| Cd5 | Atp1b1 |
| Cd63 | Il2ra |
| Sdc3 | Cd93 |
| Fam20c | Cyr61 |
| Speg | Clec4b1 |
| Serpine2 | Klra2 |
| Pdcd1 | Plbd1 |
| Dpp7 | Rasgrp4 |
| Gnas | Plekhb1 |
| Dkk2 | Cttn |
| Atp6v0d2 | Cd209a |
| Rragd | Adgre4 |
| Tnfsf8 | Trim59 |
| Fabp3 | Hid1 |
| Yipf7 | Tmem98 |
| Spp1 | F13a1 |
| Gpnmb | Ifi203 |
| Usp18 | Tbc1d10c |
| Trpm1 | Tlr8 |
| Cox6a2 | Mgl2 |
| Fgf13 | S100a10 |
| St14 | Flt3 |
| Cadm1 | Klri2 |
| Myo1e | Hepacam2 |
| Trib3 | Fpr1 |
| Slc16a8 | Clec4a1 |
| Clec1a | Cdc42ep1 |
| Mamdc2 | Vstm4 |
| Chst2 | Gja1 |
| Rtp4 | Gprc5c |

| | |
|---|---|
| Lgals3bp | Treml4 |
| Cacna1a | Fpr2 |
| Egln3 | Plekhg3 |
| Etl4 | Clec12a |
| Asb10 | Spock1 |
| Lgi2 | Ms4a4b |
| AF529169 | Cxcl12 |
| Plekhh2 | Nnat |
| Ptchd1 | Sirpb1c |
| Kcnj2 | Mndal |
| Alkbh2 | 4930523C07Rik |
| Dact1 | Sirpb1b |
| Hcar2 | Gm21188 |
| Prr15 | Ms4a4a |
| Ifit2 | Sox2ot |
| Mrgpre | AC238811.2 |
| Ch25h | |
| Lrguk | |
| Tcim | |
| Actr3b | |
| Arhgap24 | |
| Cd200r4 | |
| Ifit3b | |
| Sox7 | |
| Cst7 | |
| Gm1673 | |
| Fam189a2 | |
| Il4i1 | |
| Gas2l3 | |
| Ifit3 | |
| Tmem92 | |
| Gm11545 | |
| Serpinb1c | |
| Clec7a | |
| Gm15753 | |
| Gm13391 | |
| F630206G17Rik | |
| Gm13657 | |
| Olfr111 | |
| Olfr110 | |
| 4930556M19Rik | |
| Gm26902 | |
| Gm26714 | |
| Gm26728 | |
| Gm38335 | |
| Gm43351 | |
| AC102815.1 | |
| AC130671.2 | |
| AC130671.3 | |
| Efnb3 | |

Htra1
Rnd3
Tslp
Cyr61
Mmd2
Ank3
Ccdc162

| | logFC | AveExpr | t | P.Value | adj.P.Val |
|---|---|---|---|---|---|
| CE(18:1) | -0,45942985 | -3,0485914 | -0,91952665 | 0,2347978 | 0,447385 |
| Cer(d18:1/16:( | 0,86819798 | -4,70278086 | 3,79947516 | 0,00048499 | 0,00379909 |
| Cer(d18:1/18:( | -1,41718339 | -4,59941139 | -6,18225861 | 1,47E-06 | 3,67E-05 |
| Cer(d18:1/24:( | 0,88952242 | -4,984679 | 2,14855501 | 0,0251418 | 0,09328931 |
| Cer(d18:1/24:: | 1,08236357 | -3,65188067 | 4,04797948 | 0,00027108 | 0,00238893 |
| Cholesterol | -1,17998423 | -4,81996494 | -5,18748645 | 1,59E-05 | 0,00020359 |
| DG(16:0_18:1 | 0,43928557 | -5,74002867 | 0,83062493 | 0,270658 | 0,49799533 |
| DG(16:0_20:4 | 1,75736993 | -5,44878668 | 6,18665116 | 1,56E-06 | 3,67E-05 |
| DG(18:0_18:1 | -0,64903273 | -7,76107579 | -0,9979693 | 0,23391173 | 0,447385 |
| DG(18:0_20:4 | -0,06689945 | -5,04697047 | 0 | 0,87298619 | 0,92352186 |
| DG(18:0_22:6 | 1,56129309 | -7,93113166 | 2,14180881 | 0,03059038 | 0,11059601 |
| DG(18:1/18:1) | -0,14960969 | -5,51009057 | -0,03376452 | 0,70251323 | 0,81192103 |
| DG(18:1_20:4 | 1,32164843 | -6,36168659 | 4,46466723 | 9,98E-05 | 0,00100483 |
| HexCer(d18:1 | 0,28383999 | -8,58265014 | 0,64055448 | 0,30325358 | 0,53527129 |
| HexCer(d18:1 | -1,31833729 | -8,24957288 | -4,24548468 | 0,00017378 | 0,00163349 |
| HexCer(d18:1 | -0,41378419 | -7,11854122 | -1,45881797 | 0,08313526 | 0,20932272 |
| HexCer(d18:1 | -0,9247378 | -7,63393724 | -3,21344142 | 0,00208771 | 0,01132184 |
| HexCer(d18:1 | -2,66930309 | -6,30983969 | -6,01682489 | 2,63E-06 | 5,30E-05 |
| LPC(16:0) | -0,35710278 | -4,93599225 | -1,12503686 | 0,14550129 | 0,32564574 |
| LPC(16:1) | 0,30895087 | -9,23275481 | 0,97645013 | 0,17872353 | 0,38651339 |
| LPC(18:0) | -0,4693947 | -4,84204024 | -1,63473342 | 0,06126519 | 0,1762937 |
| LPC(18:1) | -0,11596769 | -5,11088991 | 0 | 0,59999646 | 0,73564783 |
| LPC(20:4) | 0,4483579 | -7,59248541 | 1,57470512 | 0,06806608 | 0,17772809 |
| LPC(22:6) | 0,50765707 | -10,17776 | 1,57204232 | 0,0708145 | 0,18154262 |
| LPC(26:0) | 0,41316212 | -13,1508533 | 0,7479167 | 0,30568307 | 0,53527129 |
| PC(16:0/5:0(C | -0,42430056 | -13,9160494 | -0,521748 | 0,46230487 | 0,63286395 |
| PC(16:0/9:0(C | -1,14474798 | -12,9345523 | -1,99050943 | 0,03879836 | 0,12722252 |
| PC(16:0/9:0(C | -0,53931579 | -12,4308219 | -0,7462 | 0,34250272 | 0,54878276 |
| PC(34:1) | 0,20296515 | -3,05649617 | 0,45495655 | 0,34018505 | 0,54878276 |
| PC(36:1) | -1,47891116 | -6,05254872 | -6,4185489 | 8,45E-07 | 2,98E-05 |
| PC(36:2) | 0,63373744 | -5,77519092 | 3,13687646 | 0,00237154 | 0,01238471 |
| PC(36:4) | 1,08975892 | -1,74201818 | 5,60963151 | 5,61E-06 | 9,88E-05 |
| PC(38:4) | 0,43463075 | -4,87157725 | 1,57568977 | 0,06745314 | 0,17772809 |
| PC(38:6) | 0,31254779 | -6,48782828 | 1,14633769 | 0,13545322 | 0,31309679 |
| PC(40:5) | -0,02152846 | -10,0898335 | 0 | 0,92624957 | 0,95329335 |
| PC(40:6) | -0,60149005 | -7,67033921 | -3,24641582 | 0,00181841 | 0,01068317 |
| PC(O-16:0/0:( | 0,25090583 | -8,91527106 | 0,58693386 | 0,30974466 | 0,53527129 |
| PC(O-18:0/2:( | -0,25508909 | -11,6124151 | -0,47332947 | 0,3842502 | 0,5763753 |
| PE(34:1) | 0,28830129 | -4,29121501 | 0,70796112 | 0,2719549 | 0,49799533 |
| PE(36:1) | -1,2194381 | -4,32665617 | -5,31891493 | 1,15E-05 | 0,00016277 |
| PE(36:2) | -0,60274812 | -4,37525016 | -1,62229293 | 0,06779612 | 0,17772809 |
| PE(36:4) | 1,53607473 | -3,66653247 | 9,10749455 | 2,54E-09 | 3,58E-07 |
| PE(38:4) | 0,15461697 | -2,06593151 | 0,11428043 | 0,48678794 | 0,65355571 |
| PE(38:5) | 0,62419517 | -4,15770664 | 2,92756573 | 0,00388843 | 0,01958103 |
| PE(38:6) | 0,53620912 | -2,38259038 | 2,86657585 | 0,004435 | 0,02156326 |
| PE(40:4) | -0,79143217 | -4,85388851 | -3,97175487 | 0,00031338 | 0,00259918 |

| | | | | | |
|---|---|---|---|---|---|
| PE(40:6) | -0,48005907 | -2,37357847 | -2,46682363 | 0,01090951 | 0,04644083 |
| PE(40:7) | 0,78987081 | -3,59676799 | 4,54179791 | 7,59E-05 | 0,00082349 |
| SM(d18:1/16:( | -0,34706747 | -4,26913244 | -1,88248954 | 0,03645201 | 0,1223746 |
| SM(d18:1/18:( | -0,64871651 | -6,76981832 | -3,47204661 | 0,00105813 | 0,00678166 |
| SM(d18:1/24:( | -0,12810309 | -8,23981688 | 0 | 0,61726881 | 0,75030088 |
| SM(d18:1/24:1 | -0,14112266 | -5,86454379 | -0,02693636 | 0,51419797 | 0,66422338 |
| TG(18:0_36:2 | 0,38975274 | -11,4567928 | 1,02131338 | 0,18092116 | 0,38651339 |
| TG(18:1_34:2 | 0,3313046 | -11,1133461 | 0,66731329 | 0,31591375 | 0,53527129 |
| TG(18:1_34:3 | 0,40346641 | -14,2797004 | 0,74748899 | 0,30245435 | 0,53527129 |
| TG(20:4_32:1 | 0,60950856 | -14,9379551 | 0,97445029 | 0,23861629 | 0,44859862 |
| TG(20:4_34:2 | 0,14647435 | -15,3715926 | 0,02735224 | 0,68707719 | 0,80211311 |
| TG(20:4_36:2 | 0,46351324 | -14,6006493 | 0,99198113 | 0,20627068 | 0,40394676 |
| 1-Methylnicoti | -0,50464177 | -10,3560462 | -0,59511373 | 0,43336792 | 0,61612604 |
| 3-Hydroxykynu | 0,36222126 | -11,540233 | 0,25529722 | 0,6883382 | 0,80211311 |
| 5'-Methylthioa | 1,87447151 | -9,80036875 | 3,33011964 | 0,00190481 | 0,01074312 |
| Acetylcarnitine | -0,81593732 | -9,5357 | -1,97116234 | 0,03610125 | 0,1223746 |
| Adenosine | 3,38031779 | -6,92534652 | 7,29006283 | 1,49E-07 | 1,05E-05 |
| alpha-Tocophe | 0,21777748 | -12,7632506 | 0,26534191 | 0,52286524 | 0,66422338 |
| Arabitol | 0,03712196 | -2,45764416 | 0 | 0,87395902 | 0,92352186 |
| Arginine | 0,17131243 | -8,91940746 | 0,15582666 | 0,52289925 | 0,66422338 |
| Aspartic acid | 0,230186 | -6,51264938 | 0,31376878 | 0,49132557 | 0,65355571 |
| Asymmetric di | 0,30653672 | -3,52234555 | 0,63776077 | 0,31888503 | 0,53527129 |
| Butyrobetaine | 0,0877062 | -11,8560353 | 0 | 0,81471978 | 0,89050767 |
| Butyrylcarnitin | -1,02720211 | -11,3812032 | -3,72038142 | 0,00060975 | 0,00452495 |
| Carnitine | -0,43819065 | -9,75685318 | -1,14566435 | 0,15142572 | 0,33360979 |
| Choline | 1,16539584 | -6,30361673 | 2,53959461 | 0,0111985 | 0,04644083 |
| Creatine | 1,85907778 | -2,19028661 | 3,64840757 | 0,00085644 | 0,00603789 |
| Curcumin | -0,44899854 | -11,3177167 | -1,00533428 | 0,19839732 | 0,3940003 |
| Cysteinylglycir | 0,20389602 | -5,46691167 | 0,3429822 | 0,41301542 | 0,6066164 |
| Decanoylcarni | -0,01881338 | -12,8097428 | 0 | 0,95039735 | 0,9571859 |
| Dimethylethan | 0,10709513 | -9,28184695 | 0 | 0,65571895 | 0,78352858 |
| Dopamine | -0,0155734 | -9,61843035 | 0 | 0,94829977 | 0,9571859 |
| Glucose | 0,48264586 | -6,7541921 | 1,66398533 | 0,05824229 | 0,17338361 |
| Glutamic acid | 0,38434416 | -3,95082562 | 0,97629978 | 0,19500942 | 0,3940003 |
| Glutamine | -0,26565957 | -6,86837202 | -0,44062978 | 0,42150705 | 0,61270612 |
| Glutathione | 1,04468358 | -4,46001186 | 2,44132188 | 0,01360308 | 0,05480098 |
| Glycerophospl | 0,79625119 | 3,01959034 | 1,38758799 | 0,1202875 | 0,29242307 |
| Hexanoylcarni | 0,48738764 | -10,6511544 | 0,7246289 | 0,34246609 | 0,54878276 |
| Histidine | -0,01296987 | -9,52843995 | 0 | 0,97202562 | 0,97202562 |
| Homocysteine | 0,78415023 | -1,78699341 | 1,7343702 | 0,05902421 | 0,17338361 |
| Homoserine | -0,4927004 | -5,91442892 | -1,05502133 | 0,18848108 | 0,39082107 |
| Hydroxybutyry | 0,47394882 | -8,49658383 | 0,65669309 | 0,38160471 | 0,5763753 |
| Hydroxyisovale | 0,2545489 | -11,2858422 | 0,27692369 | 0,57393565 | 0,7161498 |
| Imidazoleaceti | -0,81978069 | -2,98930733 | -1,91029179 | 0,0412449 | 0,13217116 |
| Inosine | -0,58713796 | -13,0413808 | -0,60756589 | 0,4437561 | 0,61950109 |
| Isovalerylcarni | -0,85975792 | -13,2589222 | -2,75893375 | 0,00611891 | 0,02875888 |
| Leucine | -0,08428646 | -9,97207122 | 0 | 0,74632647 | 0,84864542 |

| | | | | | |
|---|---|---|---|---|---|
| Lysine | -0,0327599 | -12,7215209 | 0 | 0,93734945 | 0,9571859 |
| Methionine | -0,22394934 | -8,64380578 | -0,49348219 | 0,33856235 | 0,54878276 |
| N-Acetylaspar | -0,04365507 | 5,61126796 | 0 | 0,88987945 | 0,92942965 |
| N6,N6,N6-Trir | 0,26157369 | -11,7520546 | 0,42387654 | 0,4309833 | 0,61612604 |
| Niacinamide | -1,10971806 | -10,1038201 | -2,03486014 | 0,03483432 | 0,1223746 |
| Norepinephrin | -0,1416763 | -13,5973267 | -0,01169697 | 0,71641087 | 0,82125148 |
| Octanoylcarnit | -0,24736584 | -10,477005 | -0,49878164 | 0,35842427 | 0,56153135 |
| Ornithine | -0,38618488 | -9,30201781 | -0,48897324 | 0,47186215 | 0,63973618 |
| Palmitoylcarni | -1,56277046 | -10,2318053 | -6,55547264 | 6,20E-07 | 2,91E-05 |
| Phenylalanine | 0,17106709 | -9,61561482 | 0,16966239 | 0,49971662 | 0,65850508 |
| Propionylcarni | -0,85701082 | -10,3209001 | -3,26541146 | 0,00180316 | 0,01068317 |
| Pyridoxamine | -0,22758359 | -5,50779822 | -0,54141318 | 0,31611692 | 0,53527129 |
| Pyroglutamic a | -1,10985445 | -7,26010187 | -2,60145827 | 0,0095357 | 0,0420167 |
| S-Adenosylme | 1,88289824 | -10,3423853 | 4,85933647 | 4,07E-05 | 0,00047823 |
| Serine | 0,29238129 | -3,95395847 | 0,54593884 | 0,36707252 | 0,56257854 |
| Stearoylcarniti | -1,08030444 | -13,1540846 | -3,51439602 | 0,00103096 | 0,00678166 |
| Sucrose | -1,24200545 | -5,89970521 | -1,22696536 | 0,18597033 | 0,39082107 |
| Taurine | -0,51093568 | 3,23489534 | -1,35714566 | 0,10789249 | 0,26689196 |
| Trimethylamin | -0,05678645 | -9,98272145 | 0 | 0,8048017 | 0,89050767 |
| Valerobetaine | 0,14256525 | -7,35203047 | 0,0089383 | 0,8093319 | 0,89050767 |
| Xanthosine | 0,04185544 | -8,64370092 | 0 | 0,85994062 | 0,92352186 |
| Fructose 1,6-b | 0,58330619 | -3,73237864 | 1,81355096 | 0,04534099 | 0,14206844 |
| Glucose 6-pho | -0,34429634 | -8,44349782 | -0,60398806 | 0,36248774 | 0,56165683 |
| Lactic acid | -0,69699554 | -4,26880164 | -1,35016408 | 0,12329955 | 0,29466504 |
| Phosphocreat | 0,3090714 | -4,65322611 | 0,38346699 | 0,51692122 | 0,66422338 |
| Sedoheptulose | 0,23918717 | -5,46391746 | 0,24127638 | 0,59621938 | 0,73564783 |
| Succinic acid | 0,05937267 | -1,48303748 | 0 | 0,7977631 | 0,89050767 |
| LacCer(d18:1/ | -1,20035312 | -11,2229595 | -2,43026678 | 0,01459097 | 0,05714796 |
| Sitosteryl hexo | 0,82455694 | -4,09031282 | 0,8074142 | 0,34903432 | 0,55296449 |
| 3-Hydroxytyro | -0,38731961 | 1,39727676 | -0,40663958 | 0,5450508 | 0,68618002 |
| Hydroxyhexan | 0,45964752 | -12,7093842 | 0,57156901 | 0,43696882 | 0,61612604 |
| Threonine | -0,07143635 | -6,38336219 | 0 | 0,87767326 | 0,92352186 |
| Tryptophan | -0,09593653 | -13,8577941 | 0 | 0,83173817 | 0,90211602 |
| LacCer(d18:1/ | 0,05316038 | -10,5386011 | 0 | 0,90664643 | 0,93997902 |
| TG(20:4_36:0) | 0,97901866 | -15,6597464 | 2,31096637 | 0,01798002 | 0,06851846 |
| 9-Hexadecenc | -0,30000904 | -11,9687832 | -0,50342802 | 0,40417731 | 0,59988422 |
| Hypoxanthine | -0,24924269 | 1,17357087 | -0,18903391 | 0,68567519 | 0,80211311 |
| Methionine S- | 0,52143628 | -7,00101525 | 1,21661582 | 0,14236955 | 0,32377591 |
| MG(18:1) | 0,76622426 | -3,58163892 | 1,70246838 | 0,06257716 | 0,17646759 |
| TG(20:4_36:3) | 0,72508343 | -14,9889208 | 1,73685018 | 0,05711356 | 0,17338361 |
| Ergothioneine | -5,03087043 | -8,8452916 | -5,48674302 | 1,13E-05 | 0,00016277 |
| gamma-Aminc | 0,197693 | -15,0202396 | 0,1488125 | 0,64950553 | 0,78273743 |
| Sarcosine | 0,46170374 | -8,83844572 | 1,013665 | 0,19781435 | 0,3940003 |
| Thiamine | -0,33059449 | -12,9350895 | -0,47728679 | 0,44859642 | 0,62011858 |
| Ethanolamine | -0,11032449 | -9,49043629 | 0 | 0,78766702 | 0,8884884 |
| Coenzyme Q1 | 1,70822743 | -9,1267996 | 2,66219043 | 0,00945294 | 0,0420167 |
| Alanine | 0,54633486 | -8,99918137 | 1,61542449 | 0,06650396 | 0,17772809 |

| | | | | | |
|---|---|---|---|---|---|
| Histamine | 0,80495339 | -4,43650813 | 1,35547739 | 0,12860834 | 0,30222961 |

| | logFC | AveExpr | t | P.Value | adj.P.Val |
|---|---|---|---|---|---|
| FDG-MP | -0,20889435 | -7,22607002 | 0 | 0,6953915 | 0,84238482 |
| Sum of Citric a | -1,19771405 | -0,71335699 | -2,18455292 | 0,01920315 | 0,06087809 |
| Glucose 6-pho | -0,0938689 | -2,06327935 | 0 | 0,86504319 | 0,9408134 |
| Lactic acid | -1,02825786 | -1,26912393 | -2,09540276 | 0,02284796 | 0,07072959 |
| Ribose 5-phos | 0,41457058 | -3,75246499 | 0,6575017 | 0,26093053 | 0,43683875 |
| Succinic acid | 0,27765064 | 2,61213553 | 0,03696857 | 0,57601613 | 0,73988278 |
| 8-Hydroxygua | -0,76624069 | -3,33580485 | -1,83058023 | 0,03876408 | 0,1099353 |
| Acetylcarnitine | -0,17497999 | -8,89738395 | 0 | 0,68944239 | 0,84202391 |
| Adenosine | -0,25924218 | -5,86660176 | 0 | 0,52934577 | 0,73712635 |
| Alanine | 0,09242368 | -4,04194929 | 0 | 0,83241121 | 0,92559156 |
| Arabitol | 0,12341892 | -5,77239699 | 0 | 0,89771514 | 0,96229897 |
| Arginine | -0,85587662 | -5,84590927 | -2,35119758 | 0,01267698 | 0,04497308 |
| Asymmetric di | -0,69255638 | -0,56328404 | -1,55510268 | 0,06583256 | 0,15490649 |
| Carnitine | -0,70520321 | -6,83658994 | -2,33417492 | 0,01312773 | 0,0454891 |
| Choline | 0,13690042 | -6,92089606 | 0 | 0,78667307 | 0,91568038 |
| Creatine | -0,38364476 | 0,24649 | -0,65685405 | 0,25873567 | 0,43683875 |
| Creatinine | -1,85403112 | -1,28124343 | -3,2937741 | 0,00130062 | 0,00842575 |
| Cysteinylglycin | -0,33020489 | -5,89682599 | -0,5513414 | 0,29267688 | 0,4740093 |
| Dimethylethan | -0,94771137 | -1,1861563 | -1,43850743 | 0,08823763 | 0,19054214 |
| Glucose | 0,53566272 | -1,7936998 | 1,7006249 | 0,0495042 | 0,12717457 |
| Glutamic acid | -0,105755 | -0,80306609 | 0 | 0,84325825 | 0,93070725 |
| Glutamine | 0,2432185 | 5,6635273 | 0 | 0,56057713 | 0,73988278 |
| Glycerophospl | -0,83546009 | 5,72149451 | -3,18445289 | 0,00164769 | 0,00982025 |
| Histidine | -1,03841002 | -5,837732 | -2,08798489 | 0,02326007 | 0,07072959 |
| Homocysteine | 0,12862707 | -3,15795248 | 0 | 0,94413117 | 0,9769135 |
| Homoserine | -0,40073382 | -0,68973218 | -0,47474388 | 0,33368953 | 0,51124396 |
| Imidazoleacet | -0,18074868 | -1,44590958 | 0 | 0,74949427 | 0,90060199 |
| Kynurenic acic | -0,44062887 | -5,82295563 | -0,5192563 | 0,32770471 | 0,50862502 |
| Mannose | 0,33998852 | -2,96850213 | 0,5304069 | 0,29990743 | 0,47538518 |
| Niacinamide | -0,46393372 | -10,018188 | -0,70193796 | 0,25213276 | 0,43683875 |
| Phenylalanine | -0,54171072 | -6,32853233 | -1,32831537 | 0,09717135 | 0,20683616 |
| Proline | -0,75797825 | -3,89992319 | -1,85254126 | 0,0370428 | 0,10822307 |
| Propionylcarni | -0,50980823 | -7,69105643 | -0,89610797 | 0,19283959 | 0,37315713 |
| Pyridoxamine | 0,0265603 | -3,00153159 | 0 | 0,95193613 | 0,97819644 |
| Pyroglutamic a | -0,47979402 | -1,25286553 | -0,37543902 | 0,45882006 | 0,65734797 |
| Serine | -0,11655433 | -0,15916651 | 0 | 0,81504422 | 0,92559156 |
| Sucrose | -1,03321629 | -3,37042588 | -1,7000289 | 0,0533139 | 0,13239619 |
| Taurine | 0,13408418 | 2,73444407 | 0 | 0,79277026 | 0,91568038 |
| Threonine | -0,56955102 | -2,00648339 | -0,85606292 | 0,21265032 | 0,40107465 |
| Trimethylamin | 0,02532638 | -3,32769598 | 0 | 0,9987234 | 0,9987234 |
| Xanthine | -0,43409325 | -4,61864935 | -0,66919613 | 0,25936973 | 0,43683875 |
| Cer(d18:1/18:( | -0,68927339 | -1,02308287 | -1,65493991 | 0,05443227 | 0,13295751 |
| Cer(d18:1/24:( | -0,24859328 | -2,34492007 | 0 | 0,56162995 | 0,73988278 |
| DG(16:0_18:1 | -0,0489107 | -0,7820098 | 0 | 0,9376189 | 0,9769135 |
| DG(18:0_22:6 | -1,58210749 | -1,98570877 | -1,47600256 | 0,0987152 | 0,20716289 |
| DG(18:1/18:1) | -0,19476891 | 0,72761201 | 0 | 0,6731283 | 0,83130935 |

| | | | | | |
|---|---|---|---|---|---|
| LPC(16:0) | -0,62881835 | -4,92066124 | -2,7211213 | 0,00527066 | 0,02181467 |
| LPC(18:0) | 0,26821384 | -2,79266994 | 0,02845599 | 0,49198566 | 0,69815108 |
| LPC(18:1) | -0,26320058 | -5,26526015 | -0,0017766 | 0,49929863 | 0,7018443 |
| MG(16:0) | -0,00778569 | 4,24095936 | 0 | 0,99030375 | 0,99699499 |
| MG(18:0) | 0,12016367 | 6,60941346 | 0 | 0,86047453 | 0,9408134 |
| MG(18:1) | 0,0214411 | 0,80445425 | 0 | 0,97591348 | 0,98919121 |
| PC(16:0/5:0(C | -1,13666814 | -6,68680335 | -2,70891055 | 0,0055159 | 0,02218794 |
| PC(16:0/9:0(C | -1,53690556 | -4,65547741 | -4,11392679 | 0,00013393 | 0,00124719 |
| PC(16:0/9:0(C | -0,96071247 | -6,60008115 | -2,81821481 | 0,00417962 | 0,01887163 |
| PC(34:1) | -0,81373728 | -0,05922635 | -5,08325192 | 8,30E-06 | 0,0004122 |
| PC(36:1) | -1,06275909 | -2,51956961 | -6,40443789 | 1,89E-07 | 1,41E-05 |
| PC(36:2) | -0,54761678 | -2,42475485 | -2,96496495 | 0,00287757 | 0,01531277 |
| PC(36:4) | -0,5857965 | -1,14395013 | -2,40959967 | 0,01102502 | 0,04006654 |
| PC(38:4) | -0,71954758 | -2,56599537 | -4,39467138 | 5,96E-05 | 0,00113529 |
| PC(38:6) | -0,82626949 | -3,72299022 | -2,89377681 | 0,00345596 | 0,0170297 |
| PC(40:5) | -0,50494316 | -7,05705899 | -1,72416867 | 0,0472848 | 0,12581134 |
| PC(40:6) | -0,91443858 | -5,07986674 | -3,33109544 | 0,00112301 | 0,00760586 |
| PE(36:2) | -0,8621439 | -0,88741279 | -4,33842775 | 6,99E-05 | 0,00113529 |
| PE(38:4) | -1,00829537 | 0,47888914 | -4,33030665 | 7,21E-05 | 0,00113529 |
| PE(40:6) | -1,18011836 | 0,58768188 | -4,20283726 | 0,00010368 | 0,00118832 |
| SM(d18:1/16:( | -0,21660633 | -2,08073505 | 0 | 0,6231748 | 0,7802777 |
| SM(d18:1/18:( | -1,05907468 | -2,69098573 | -6,67534416 | 8,80E-08 | 1,31E-05 |
| SM(d18:1/24:( | -1,38760508 | -4,63529592 | -2,89403099 | 0,00354309 | 0,0170297 |
| SM(d18:1/24:1 | -0,87541338 | -3,24263737 | -4,22358499 | 9,66E-05 | 0,00118832 |
| TG(18:0_36:2 | 0,11390005 | -0,11007115 | 0 | 0,90824591 | 0,96663315 |
| TG(18:1_34:2 | 0,09495702 | 1,55132257 | 0 | 0,92581031 | 0,97144884 |
| TG(18:1_34:3 | 0,17853932 | -1,17280716 | 0 | 0,78112896 | 0,91568038 |
| TG(20:4_34:2 | -0,08491216 | -4,48610671 | 0 | 0,89249167 | 0,96229897 |
| Cholesterol su | -0,61002692 | 0,02658479 | -0,49268503 | 0,42505782 | 0,62091779 |
| LPE(18:0) | -0,77074187 | -2,65197604 | -2,22018799 | 0,01696644 | 0,0549565 |
| PS(18:0_22:6) | -0,80788495 | 0,55717615 | -3,36693222 | 0,00101811 | 0,0075849 |
| Butyrylcarnitin | -0,43338055 | -9,63345753 | -0,7321201 | 0,23748705 | 0,43153134 |
| Ethanolamine | 0,27418405 | -6,97737734 | 0,02796908 | 0,582705 | 0,74207731 |
| Cer(d18:1/16:( | -0,11596943 | -2,05348764 | 0 | 0,82020392 | 0,92559156 |
| Cer(d18:1/24:1 | -0,58615938 | -1,80002683 | -1,39635763 | 0,08671436 | 0,19000646 |
| DG(18:0_20:4 | -0,24831367 | -3,04618152 | 0 | 0,56857157 | 0,73988278 |
| HexCer(d18:1 | -1,26670014 | -2,45934407 | -4,14623303 | 0,00012164 | 0,00124719 |
| PE(38:6) | -0,86348779 | -0,70445165 | -3,15222734 | 0,00179138 | 0,01026599 |
| PI(18:0_20:4) | -0,8532966 | -0,98403965 | -4,86688032 | 1,55E-05 | 0,00054283 |
| Glutathione | 0,04340762 | -3,54975143 | 0 | 0,92325189 | 0,97144884 |
| Tryptophan | -0,12478518 | -9,13628197 | 0 | 0,77474754 | 0,91568038 |
| HexCer(d18:1 | -1,18282478 | -4,12946809 | -4,12156418 | 0,00013023 | 0,00124719 |
| LPC(20:4) | -0,44649778 | -7,23590717 | -1,08712849 | 0,14277472 | 0,28535398 |
| PE(34:1) | -0,92579529 | -0,69638116 | -3,52328513 | 0,00067317 | 0,00527909 |
| PE(36:1) | -1,11259217 | -0,83352876 | -3,6274008 | 0,00050899 | 0,0042133 |
| PE(38:5) | -0,68806958 | -1,2266661 | -1,81472649 | 0,03977935 | 0,1099353 |
| PE(P-18:0/20: | -0,77233237 | 0,74352475 | -2,94475165 | 0,00303752 | 0,01560655 |

| | | | | | |
|---|---|---|---|---|---|
| Valerobetaine | -0,56807208 | -5,14875938 | -0,75032934 | 0,25410054 | 0,43683875 |
| PE(36:4) | -0,62322365 | -0,68561153 | -1,54520133 | 0,06653702 | 0,15490649 |
| PE(40:4) | -0,76168259 | -1,58484456 | -2,27413625 | 0,01504214 | 0,05093817 |
| (3-O-sulfo)Gal | -0,98713151 | -0,69456033 | -2,6951863 | 0,00565867 | 0,02218794 |
| PE(P-18:0/22: | -0,84076836 | -1,59011936 | -2,77219916 | 0,00467113 | 0,01988569 |
| PS(18:0_18:1) | -1,10307961 | 0,39357741 | -4,81389056 | 1,82E-05 | 0,00054283 |
| 5'-Methylthioa | -0,35442987 | -6,87472284 | -0,31135095 | 0,40062546 | 0,59102172 |
| Inosine | -0,81122769 | -6,30949874 | -1,74018883 | 0,04678191 | 0,12581134 |
| HexCer(d18:1 | -0,95972903 | -4,35366791 | -3,33628161 | 0,00110811 | 0,00760586 |
| HexCer(d18:1 | -1,28713145 | -3,47586075 | -4,3099632 | 7,67E-05 | 0,00113529 |
| LPC(22:6) | -0,41338818 | -8,87301763 | -0,60988369 | 0,27819357 | 0,45632603 |
| PC(O-16:0/2:0 | -0,79906118 | -8,20689822 | -1,72317611 | 0,04830846 | 0,12628002 |
| PE(40:7) | -1,01969041 | -1,65835709 | -3,67875788 | 0,00044228 | 0,00387644 |
| (3-O-sulfo)Gal | -1,09127806 | -0,42377043 | -4,43195926 | 5,43E-05 | 0,00113529 |
| (3-O-sulfo)Gal | -0,80706141 | 1,56801473 | -1,48662411 | 0,07680688 | 0,17339735 |
| PE(O-18:0/20: | -0,67123664 | -1,93238052 | -1,81349253 | 0,03984232 | 0,1099353 |
| PE(P-16:0/20: | -0,5092336 | 1,00441349 | -1,20425096 | 0,11913962 | 0,24317539 |
| PE(P-16:0/22: | -0,67570486 | 0,51549565 | -2,05991265 | 0,02396843 | 0,07142594 |
| PE(P-16:0/22: | -0,80822828 | -1,48227679 | -2,77452188 | 0,00464298 | 0,01988569 |
| PE(P-18:0/18: | -1,1440886 | 0,19923332 | -3,20913332 | 0,00155243 | 0,00963801 |
| PE(P-18:1/20: | -0,7726557 | 0,57910877 | -2,84112846 | 0,00393854 | 0,01833884 |
| PE(P-18:1/22: | -1,00601631 | -2,61778864 | -4,27714976 | 8,38E-05 | 0,00113529 |
| PG(16:0_18:1 | -0,77610158 | -5,03424681 | -2,49656267 | 0,00904058 | 0,03367615 |
| PI(16:0_18:1) | -0,60982433 | -4,51699121 | -1,4468717 | 0,07947207 | 0,17673639 |
| PI(16:0_20:4) | -0,56329266 | -2,17098447 | -1,51568856 | 0,06997637 | 0,16040737 |
| PS(16:0_18:1) | -0,33450784 | -2,02076099 | -0,32030568 | 0,38131303 | 0,56815641 |
| PS(18:0_20:4) | -0,58690499 | 0,28853215 | -1,68414678 | 0,05115347 | 0,12918419 |
| PS(18:1/18:1) | -0,50903515 | -1,56533512 | -1,26200243 | 0,1083824 | 0,22429135 |
| Thiamine | -0,94680092 | -10,533407 | -2,56920578 | 0,00765187 | 0,02923406 |
| (3-O-sulfo)Gal | -1,05020032 | -0,45445792 | -3,05252752 | 0,00232144 | 0,01281093 |
| Oleic acid | -0,48306574 | 0,06869625 | -0,6956611 | 0,25831556 | 0,43683875 |
| Palmitic acid | -0,3499848 | 3,59396648 | -0,42703706 | 0,33872448 | 0,51124396 |
| PI(18:1/18:1) | -0,46566334 | -4,36866471 | -0,82159812 | 0,21175392 | 0,40107465 |
| Hydroxyisoval | -0,36574632 | -8,99318556 | -0,46812601 | 0,32518784 | 0,50862502 |
| DG(16:0_20:4 | 0,22226017 | -3,43744758 | 0 | 0,60439469 | 0,76317634 |
| BMP(18:1/18:1 | -0,53604981 | -3,19088371 | -1,08187837 | 0,14554968 | 0,28535398 |
| BMP(22:6/22:6 | -0,62593389 | -4,33945107 | -1,55464114 | 0,06539989 | 0,15490649 |
| PI(18:0_18:1) | -0,2437412 | -4,80756606 | 0 | 0,55586627 | 0,73988278 |
| PS(16:0_22:6) | -0,3191084 | -4,0052715 | -0,18805873 | 0,45601457 | 0,65734797 |
| (3-O-sulfo)Gal | -0,41605284 | -1,45544479 | -0,66376164 | 0,2589062 | 0,43683875 |
| GM3(d36:1) | 0,48970246 | -2,1047301 | 0,80319132 | 0,22001099 | 0,40977046 |
| PA(16:0_18:1) | -0,39597839 | -2,85359963 | -0,60153669 | 0,27869576 | 0,45632603 |
| PE(O-16:0/20: | -0,12131353 | -1,62725587 | 0 | 0,78820738 | 0,91568038 |
| Adenosine mo | 0,13196174 | -6,76423052 | 0 | 0,83089525 | 0,92559156 |
| (3-O-sulfo)Gal | -0,24061754 | -1,82423547 | 0 | 0,57268361 | 0,73988278 |
| BMP(20:4/20:4 | -0,47430896 | -3,30951977 | -0,78010037 | 0,22589121 | 0,41552827 |
| PS(16:0_20:4) | -0,39535118 | -3,21557857 | -0,55138943 | 0,29766363 | 0,47538518 |

| | | | | | |
|---|---|---|---|---|---|
| LPI(18:0) | 0,18012395 | -4,97160345 | 0 | 0,67509015 | 0,83130935 |
| PE(O-16:0/22: | -0,01945993 | -3,82129245 | 0 | 0,96573112 | 0,98557491 |
| PG(18:0_18:1 | -0,2488477 | -7,18296333 | 0 | 0,55043143 | 0,73988278 |
| PA(18:1/18:1) | -0,09906855 | -3,84828078 | 0 | 0,82143696 | 0,92559156 |
| PE(O-18:0/22: | -0,24908238 | -3,64253797 | 0 | 0,56034817 | 0,73988278 |
| PI(18:0_22:6) | 0,33605313 | -5,08244804 | 0,41954987 | 0,33968558 | 0,51124396 |
| PI(20:4/20:4) | -0,62618833 | -5,59173998 | -1,10255789 | 0,14491948 | 0,28535398 |
| Hydroxybutyry | -0,24847945 | -9,42812854 | 0 | 0,55071155 | 0,73988278 |
| N6,N6,N6-Trir | -1,08753995 | -10,2943569 | -2,26844228 | 0,0156063 | 0,05167421 |

Antibodies used

| Antibody | Company | species | dilution |
|---|---|---|---|
| mKate2 | Evrogen | rabbit | 1:500 |
| Trem2 | R&D | sheep | 1:200 |
| Iba-1 | Novusbio | goat | 1:300 |
| Aß 1-40 | Cell Signal Technology | mouse | 1:1000 |
| Alexa647 | Invitrogen | rabbit | 1:500 |
| Alexa555 | Invitrogen | mouse | 1:500 |
| Alexa488 | Invitrogen | goat | 1:500 |
| Alexa488 | Invitrogen | sheep | 1:500 |
